# Harmine Plus Exendin-4 Enhances Remission of Recent-Onset Type 1 Diabetes Following Anti-CD3 Therapy

**DOI:** 10.64898/2026.09.21.753317

**Authors:** Geming Lu, Randy Kang, Miguel Varela, Eunjin Oh, Yansui Li, Jungeun Lee, Leah Kebrom, Juan Aldaco, Jiamin Zhang, Madeline Wichman, Mingpei Li, Fouad Kandeel, Meirigeng Qi, Peng Wang, Robert J. DeVita, Alberto Pugliese, Debbie C. Thurmond, Andrew F. Stewart, Adolfo Garcia-Ocana

**Affiliations:** Department of Molecular & Cellular Endocrinology, Arthur Riggs Diabetes & Metabolism Research Institute, Beckman Research Institute, City of Hope, Duarte, CA, 91010, USA; Diabetes, Obesity and Metabolism Institute, and Division of Endocrinology, Diabetes and Bone Diseases, Icahn School of Medicine at Mount Sinai, New York, NY, 10029, USA; Department of Translational Research & Cellular Therapeutics, Arthur Riggs Diabetes & Metabolism Research Institute, Beckman Research Institute, City of Hope, Duarte, CA, 91010, USA; Department of Pharmacological Sciences, The Marie-Josée and Henry R. Kravis Drug Discovery Institute, Icahn School of Medicine at Mount Sinai, New York, NY, 10029, USA; Department of Diabetes Immunology, Arthur Riggs Diabetes & Metabolism Research Institute, Beckman Research Institute, City of Hope, Duarte, CA, 91010, USA

## Abstract

Type 1 diabetes (T1D) results from autoimmune destruction of pancreatic β-cells. While anti-CD3 therapy can delay disease progression and preserve residual β-cell function, disease reversal will likely require both immune modulation and β-cell regeneration. We found that the combination of harmine and exendin-4 (H+E) reduced inflammation-induced human β-cell apoptosis, suppressed cytokine signaling and immunogenicity pathways, and improved β-cell function. Although H+E alone did not reverse diabetes in NOD mice, low-dose anti-CD3 followed by H+E normalized blood glucose, increased insulin levels, improved glucose tolerance, expanded β-cell mass, and enhanced diabetes remission. These effects were associated with reduced pro-inflammatory T-cell responses, increased regulatory T cells, and greater expression of exhaustion-related T-cell markers, without broad lymphocyte depletion. Similar immunomodulatory effects were observed in activated human PBMCs. Transcriptomic analyses identified the lncRNA SNHG6 as a key mediator of H+E action; SNHG6 protected β-cells from cytokine-induced stress, apoptosis, and immunogenicity. Together, these findings demonstrate that H+E promotes β-cell recovery and resilience while reducing β-cell immunogenicity, enabling remission of recent-onset T1D when combined with anti-CD3 therapy. SNHG6 emerges as a novel regulator of β-cell protection during inflammation.

## Introduction

Type 1 diabetes (T1D) is a chronic autoimmune disease characterized by the progressive, selective immune-mediated destruction of pancreatic β-cells leading to the loss of endogenous insulin production (1,2). Autoreactive CD4+ and CD8+ T cells, together with innate and adaptive immune-cell networks, infiltrate pancreatic islets and promote β-cell dysfunction and death through inflammatory cytokines, cytotoxic mediators, and sustained antigen-specific immune attack (3,4). Although exogenous insulin therapy is lifesaving, there is no clinical cure for T1D that corrects the underlying autoimmune process and restores the β-cells lost. Therefore, therapeutic strategies capable of both limiting autoimmune injury and preserving and restoring functional β-cell numbers are needed to achieve durable disease modification.

Immune modulation has shown clinical benefit in T1D (5,6). Teplizumab, an anti-CD3 monoclonal antibody, is an FDA-approved immunotherapy designed to delay progression to clinical T1D in at-risk individuals and preserves endogenous insulin secretion in newly diagnosed T1D patients (6–9). Similar to most other immune therapies tested, preservation of residual insulin secretion declines over time, indicating that immune intervention alone is insufficient to provide durable disease remission (6,7). Mechanistically, anti-CD3 therapy has been associated with reduced pathogenic T-cell activity, expansion or preservation of regulatory T cells (Tregs), and induction of inhibitory and exhaustion-associated T-cell programs (10,11). These observations suggest that immune modulation may create a therapeutic window in which additional therapeutic approaches can be implemented to enhance functional β-cell recovery and resilience to reverse T1D.

Harmine (H), a small-molecule inhibitor of dual-specificity tyrosine-phosphorylation-regulated kinase 1A (DYRK1A), significantly induces human β-cell proliferation, a feature synergistically enhanced by glucagon-like peptide-1 receptor (GLP1R) agonists such as exendin-4 (E) (12–14). Based on recent studies demonstrating the capacity of this drug combination to markedly expand human β-cell numbers *in vivo* in human islet xenografts, the combination of H+E has emerged as a candidate for β-cell regenerative therapies (14). However, in autoimmune diabetes, newly formed or functionally recovered β-cells remain exposed to ongoing immune attack and the potential of the H+E drug combination to protect functional β-cells in an inflammatory environment is unknown. Therefore, there is a need to test whether H+E can restore β-cell numbers and function in the setting of autoimmune diabetes, or whether concurrent immune modulation is required.

In addition to immune-cell-mediated destruction, β-cells *per se* actively participate in T1D disease progression (15,16). Emerging evidence indicates that β-cells respond to inflammatory and metabolic stress by altering their phenotype, increasing their immunogenicity, and contributing to local inflammatory processes. Specifically, inflammatory cytokines induce β-cell stress, unfolded protein response activation, antigen-presentation pathways, chemokine production, and pro-apoptotic programs (17–19). These responses can impair insulin biosynthesis and processing, increase proinsulin release, enhance MHC-I expression, and recruit additional immune cells to the islet microenvironment (17–19). Thus, successful T1D remission may require not only suppression of autoreactive immune cells and β-cell regeneration but also recovery of stressed β-cells into a less inflamed, less immunogenic and more functionally competent state.

Long non-coding RNAs (lncRNAs) are emerging regulators of cell stress, inflammatory signaling, and cell death (20,21). Several lncRNAs have been implicated in diabetes-related tissue injury (22–24), but their roles in human β-cell responses to inflammatory stress remain incompletely understood. One of these lncRNAs, small nucleolar RNA host gene 6 (SNHG6) has been linked to cellular survival and stress responses in various disease contexts, including diabetic complications (25,26). However, SNHG6 contribution to pancreatic β-cell resilience or immunogenicity under proinflammatory conditions is unknown.

Here, we tested the potential of the pro-regenerative combination, H+E, to reduce human β-cell death and immunogenicity and to improve functional β-cell recovery in a pro-inflammatory environment. We also explored the mechanisms underlying these effects. Further, we tested the therapeutic potential of H+E to reverse early-onset T1D in NOD mice. Our findings indicate that H+E suppressed human β-cell inflammatory and immunogenicity programs, improved β-cell stress handling and insulin secretory quality, and induced the expression of SNHG6 lncRNA, a regulator of β-cell death and immunogenicity in a proinflammatory environment. We also found that pre-treatment with low-dose anti-CD3 followed by H+E administration reversed recent-onset T1D in NOD mice. This therapeutic response was associated with reduced diabetogenic T-cell activity and increased regulatory and exhaustion-associated T-cell phenotypes. These findings support a dual-compartment model in which anti-CD3 reduces autoimmune pressure while H+E promotes β-cell recovery and decreases β-cell vulnerability to inflammatory attack. Collectively, these results suggest promising therapeutic potential for the combination therapy of anti-CD3 and H+E for early-onset T1D reversal.

## Results

### Harmine plus Exendin-4 protects human β-cells from cytokine-induced inflammatory and apoptotic programs

To study the impact of H, E and H+E in the human β-cell transcriptome under inflammatory conditions, we performed single cell RNA sequencing (scRNA-seq) of isolated human islets exposed to a cocktail of pro-inflammatory cytokines (IL1β+TNFα+IFNγ) and vehicle, H, E and H+E for 6h (**Fig. 1A**). After integration, endocrine and non-endocrine islet cell populations were visualized by UMAP and annotated using conventional islet cell gene markers (**Fig. 1B-C, Suppl. Fig. 1A-E**). Pathway enrichment analysis using the UCell algorithm with escape package showed that H+E markedly suppressed apoptosis- and inflammation-related pathways induced by cytokine treatment in human β-cells beyond the suppression induced by H or E alone (**Fig. 1D**). Consistent with these transcriptomic findings, H+E, but not H or E, significantly reduced cytokine-induced human β-cell death, as assessed by TUNEL and insulin immunolabeling of human islets treated with cytokines (**Fig. 1E**). In addition, H+E also significantly reduced *CASP3* expression in cytokine-treated EndoC-βH1 cells, further supporting an anti-apoptotic effect in human β-cells (**Fig. 1F**). We next examined the effect of H+E in cytokine-induced inflammatory pathway scores in human β-cells in the scRNA-seq dataset. H+E profoundly reduced IFNγ-, IL1/IL1R-, and TNF/TNFR-related inflammatory pathway scores compared with individual drug treatments (**Fig. 1G**). In support of the suppression of inflammatory pathways, H+E significantly decreased cytokine-induced *iNOS* expression and nitric oxide (NO) production in human islets (**Fig. 1H**). Taken together, these results indicate that H+E suppresses cytokine-induced inflammatory signaling, *iNOS* expression and NO production, and apoptotic programs in human β-cells, supporting a direct β-cell protective effect under inflammatory conditions relevant to T1D.

**Figure 1.**
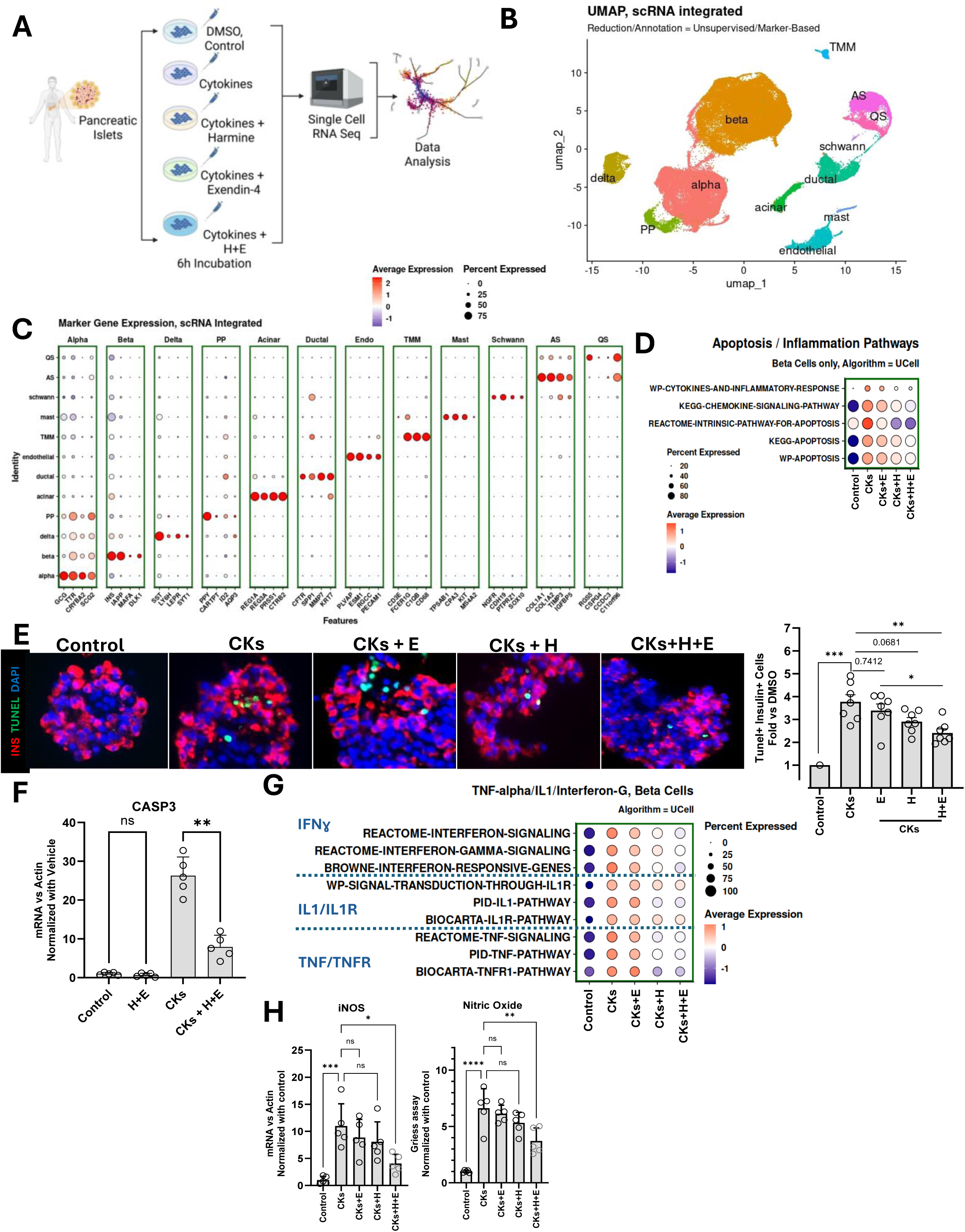
Experimental design, single cell data clustering, and apoptosis and cytokine signaling pathway analysis and validation in human islets treated with cytokines, harmine and exendin-4. **A** Human islet treatment, processing and data generation scheme. Created in BioRender. Garcia-Ocana, A. (2026). **B** UMAP with cell type annotation. PP=Pancreatic Polypeptide+ Cells; AS=Activated Stellate Cell; QS=Quiescent Stellate Cells; TMM=T Cells+Macrophages+Monocytes. **C** Dot-plot visualization of top 4 canonical genes expressed in each cluster. **D** Pathway enrichment analysis for β-cells, emphasizing apoptosis/inflammation pathways. **E** Representative images and quantitation of β-cell death by TUNEL staining of non-diabetic human islets treated with cytokines (Cks) and harmine (H) and exendin-4 (E). N=7 different human islet preparations; One-way ANOVA with Tukey’s multiple comparisons test. *P<0.05, **P<0.01. **F** *CASP3* expression by qPCR in EndoC-βH1 cells treated with Cks, H and E. N=5 experiments in duplicate. Two-tailed Student’s t-test, **P<0.01 **G** Pathway enrichment analysis for β-cells, emphasizing IL1β, TNFα and IFNγ. **H** *iNOS* expression by qPCR and Nitric Oxide levels in human islets treated with Cks, H and E. N=5 different human islet preparations; one-way ANOVA with Tukey’s multiple comparisons test, *P<0.05, **P<0.01.

### Harmine plus Exendin-4 suppresses cytokine-induced β-cell immunogenicity programs

Inflammatory cytokines not only induce β-cell death but also increase β-cell immunogenicity through IFNγ-responsive antigen-presentation and chemokine pathways (17–19). Since H+E significantly reduced cytokine-induced inflammation and apoptosis in human β-cells, we next examined whether H+E also altered β-cell immune visibility and immune-cell recruitment signals.

Analysis of the scRNA-seq dataset showed that H+E reduced cytokine-induced expression of classical MHC-I-related genes, including HLA-A, HLA-B, HLA-C, and B2M, in human β-cells compared with cytokine-treated controls and individual drug treatments (**Fig. 2A**). In contrast, expression of the immunomodulatory MHC-I molecule HLA-E was increased by cytokines and further enhanced by H+E treatment (**Fig. 2A**). Along with these transcriptomic findings, immunostaining of cytokine-treated human islet cells showed increased HLA-ABC expression in insulin-positive cells, which was markedly reduced by H+E treatment (**Fig. 2B, Suppl. Fig. 2A**). HLA-E expression was increased in insulin-positive cells by cytokines and further enhanced by H+E treatment (**Fig. 2B, Suppl. Fig. 2B**).

**Figure 2.**
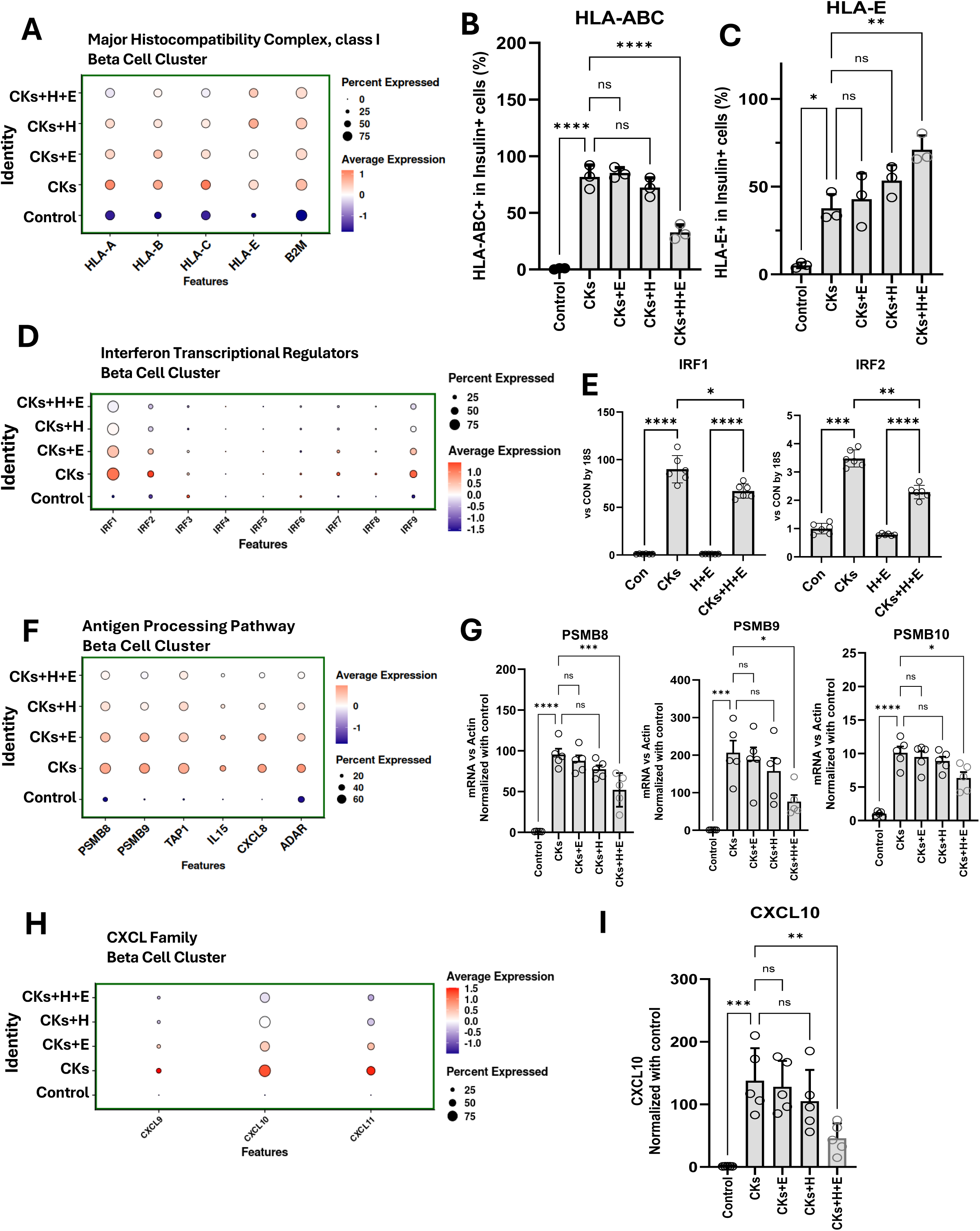
Differentially expressed genes and pathway enrichment analysis of MHC-I, IRFs, antigen presentation and chemokines in human β-cells by scRNA-seq of human islets treated with cytokines and harmine and exendin-4. **A** Expression of genes of the major histocompatibility complex, class I in β-cells of human islets treated with cytokines (Cks), harmine (H) and exendin-4 (E). **B** Quantitation of HLA-ABC and insulin immunolabeling of human islet cells treated with Cks, H and E. N=3 different human islet preparations; one-way ANOVA with Tukey’s multiple comparisons test,****P<0.0001. **C** Quantitation of HLA-E and insulin immunolabeling of human islet cells treated with Cks, H and E. N=3 different human islet preparations; one-way ANOVA with Tukey’s multiple comparisons test, *P<0.05, **P<0.01. **D** Expression of interferon regulatory factors (IRFs) transcription factor genes in β-cells. **E** Quantitation of the expression of representative IRFs, *IRF1* and *IRF2*, in human islets treated with Cks, H and E. N=6 different human islet preparations; one-way ANOVA with Tukey’s multiple comparisons test, *P<0.05, **P<0.01, ***P<0.0001. **F** Expression of genes involved in the antigen processing pathway in β-cells. **G** Quantitation of the expression of representative genes of the antigen processing pathway *PSMB8*. *PSMB9*, *PSMB10*, in human islets treated with Cks, H and E. N=5 different human islet preparations; one-way ANOVA with Tukey’s multiple comparisons test, *P<0.05, ***P<0.001, ***P<0.0001. **H** Expression of the CXCL family of genes in β-cells. **I** Quantitation of the CXCL10 levels in media of human islets treated with Cks, H and E. N=5 different human islet preparations; one-way ANOVA with Tukey’s multiple comparisons test, **P<0.01, ***P<0.001.

We next examined IFNγ-responsive transcriptional regulators. Cytokine treatment induced the expression of interferon regulatory factor family members in human β-cells, including *IRF1* and *IRF2*, whereas H+E reduced their expression compared with individual drug treatments **(Fig. 2D**). These results were confirmed by qPCR analysis where H+E decreased *IRF1* and *IRF2* expression in cytokine-treated human islets (**Fig. 2E**).

Because IFNγ signaling also regulates antigen-processing machinery (27,28), we analyzed genes involved in immunoproteasome and antigen-presentation pathways. Cytokine treatment increased expression of *PSMB8*, *PSMB9*, *TAP1*, *IL15*, *CXCL8*, and *ADAR* in human β-cells, and H+E markedly reduced the expression of these genes (**Fig. 2F**). qPCR analysis confirmed that H+E significantly decreased the expression of proteins of the immunoproteasome in cytokine-treated human islets (**Fig. 2G**).

We then examined chemokines involved in immune-cell recruitment. Cytokines increased expression of *CXCL9*, *CXCL10*, and *CXCL11* in human β-cells, whereas H+E reduced the expression of these chemokines (**Fig. 2H**). Consistent with this result, H+E treatment significantly decreased CXCL10 protein levels in the culture media of cytokine-exposed human islets compared with individual drug treatments (**Fig. 2I**). Taken together, these results indicate that H+E suppresses cytokine-induced β-cell immunogenicity programs by reducing classical MHC-I expression, IFNγ-responsive transcription factors, antigen-processing genes, and chemokine production, while enhancing HLA-E expression.

### Harmine plus Exendin-4 reduces cytokine-induced β-cell stress and rescues insulin secretion in human islets

Cytokine-induced β-cell inflammation is closely associated with cellular stress, unfolded protein response activation, and impaired insulin processing (17–19). Since H+E reduced inflammatory and immunogenicity-related programs in cytokine-treated human β-cells, we next examined whether H+E also attenuated β-cell stress responses and improved insulin secretory function.

Pathway enrichment analysis of the scRNA-seq dataset showed that cytokine treatment increased stress- and unfolded protein response-related processes in human β-cells (**Fig. 3A**). H+E reduced the expression of genes in these pathways induced by cytokine treatment compared with individual drug treatments (**Fig. 3A**). Consistent with this reduction in stress-response pathways, H+E significantly reduced human β-cell death induced by the ER-stress inducer thapsigargin (**Fig. 3B**). To further validate these findings, we measured representative stress-response gene expression in cytokine-treated human islets by qPCR. H+E significantly reduced the expression of *ATF6* and spliced *XBP1*, markers of canonical UPR signaling, as well as *ATF3*, *TXNIP*, and *DDIT3* (CHOP), which reflect integrated stress response genes, stress-associated inflammatory injury genes, and maladaptive ER-stress/apoptotic signaling genes (**Fig. 3C-F**).

**Figure 3.**
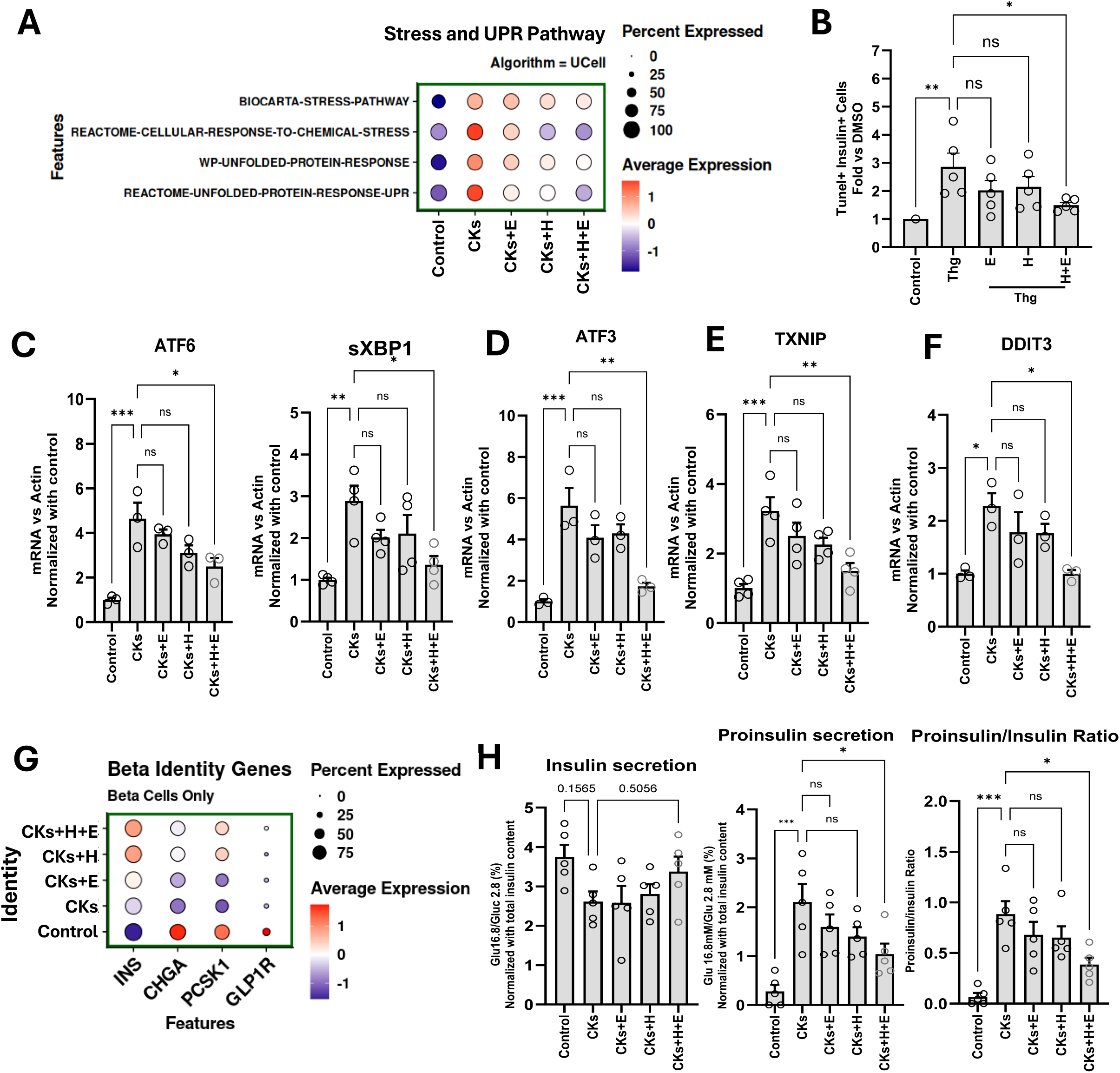
Differentially expressed genes and pathway enrichment analysis of stress and UPR, and β-cell identity and function in human β-cells by scRNA-seq of human islets treated with cytokines, harmine and exendin-4. **A** Expression of genes of the UPR-ER Stress pathway in β-cells of human islets treated with cytokines (Cks) and harmine (H) plus exendin-4 (E). **B** Quantitation of β-cell death by TUNEL staining of non-diabetic human islets treated with the ER stress inducer Thapsigargin (Thg), H and E. N=5 different human islet preparations; one-way ANOVA with Tukey’s multiple comparisons test, *P<0.05, **P<0.01. **C** *ARF6 and sXBP1*, **D** *ATF3*, **E** *TXNIP* and **F** *DDIT3* (CHOP) expression by qPCR in human islets treated with Cks, H and E. N=4 different human islet preparations; one-way ANOVA with Tukey’s multiple comparisons test, *P<0.05, **P<0.01, ***P<0.001. **G** Expression of β-cell identity genes in human islets treated with Cks, H and E. **H** Insulin and proinsulin secretion per total insulin content in human islets treated with Cks, H and E. Calculation of the proinsulin and insulin ratio in these studies. N=5 different human islet preparations; one-way ANOVA with Tukey’s multiple comparisons test, *P<0.05, ***P<0.001.

We next examined whether this reduction in β-cell stress by H+E was associated with preservation of β-cell functional gene expression. In the scRNA-seq dataset, cytokine treatment altered expression of genes involved in insulin production, proinsulin processing, secretory granule biology, and GLP-1 responsiveness, including *INS*, *CHGA*, *PCSK1*, and *GLP1R*. H+E partially restored or enhanced the expression of these genes in cytokine-treated β-cells (**Fig. 3G**). Based on these results, we next assessed insulin and proinsulin secretion in cytokine- and H+E-treated human islets. Cytokine exposure impaired insulin secretion and increased proinsulin release, resulting in a significantly elevated proinsulin/insulin ratio (**Fig. 3H**). H+E treatment partially restored insulin secretion, markedly reduced proinsulin release, and significantly decreased the proinsulin/insulin ratio (**Fig. 3H**). Collectively, these results indicate that H+E reduces cytokine-induced β-cell stress and improves insulin secretory quality in human islets exposed to inflammatory conditions relevant to T1D.

### Harmine plus Exendin-4 reverses early-onset T1D in NOD mice pre-treated with anti-CD3

*In vivo* treatment with H+E remarkably and safely enhanced human β-cell mass in a mouse model of human islet xenografts (14). Harmine enhanced Treg differentiation, impaired Th17 differentiation and attenuated inflammation in multiple experimental mouse models of systemic autoimmunity and mucosal inflammation (29). Treatment with the GLP1R agonist semaglutide, in patients soon after the diagnosis of T1D, increased C-peptide levels and improved glycemic control (30). Results in the current study indicate that H+E significantly decreases death, immunogenicity, stress and recovers function of human β-cells exposed to a proinflammatory environment (**Figs. 1-3**). Based on these collective observations, we next explored whether H+E treatment could reverse diabetes in early-onset T1D NOD mice. Diabetic female NOD mice (>250 mg/dl blood glucose for three days) were treated with continuous infusion of H, E or H+E for 56 days, changing the infusion pumps every 28 days, as previously described in detail (14). As shown in **Suppl. Figs. 3A and 3B**, none of the three treatments improved blood glucose levels or reduced the incidence of diabetes in these mice. Plasma insulin on day 56 was barely detectable and not different among the different treatment groups (**Suppl. Fig. 3C**). These studies indicate that the combination therapy at the doses tested cannot reverse diabetes in early-onset T1D NOD mice.

Clinical trials have demonstrated the efficacy and safety of anti-CD3 antibodies, such as teplizumab (Tzield), in preventing or delaying the progression of T1D, particularly in individuals at high risk of developing the disease (6–9). Furthermore, teplizumab administration in patients with newly diagnosed type 1 diabetes resulted in significantly higher stimulated C-peptide levels than in patients receiving placebo. However, other secondary endpoints such as the insulin doses that were required to achieve glycemic goals, glycated hemoglobin levels, time in the target glucose range, and clinically important hypoglycemic events did not significantly change with teplizumab treatment (6–9). Since H+E enhances β-cell mass and reverses diabetes in streptozotocin (STZ)-treated diabetic mice (12–14) but fails to do so in early-onset T1D NOD mice (**Suppl Fig. 3A-C**), we tested whether transient immune modulation with anti-CD3 could create a permissive window for H+E-mediated β-cell recovery. Recent-onset diabetic NOD mice were treated with low-dose anti-CD3 for three days (31), followed by continuous infusion of vehicle, H, E, or H+E for 56 days (14) (**Fig. 4A**). Anti-CD3+H+E rapidly reduced blood glucose and maintained glycemia below the diabetic range throughout treatment, whereas anti-CD3 followed by vehicle or either individual drug produced only partial or limited improvement (**Fig. 4B**). Consistently, anti-CD3+H+E markedly reduced the fraction of diabetic mice (**Fig. 4C**), increased circulating insulin levels (**Fig. 4D**), and improved glucose tolerance (**Fig. 4E-F**). We next examined whether improved glycemic control was associated with recovery of β-cell mass and β-cell resilience. Anti-CD3+H+E increased β-cell mass compared with control and individual treatment groups (**Fig. 4G**). This increase was associated with enhanced β-cell proliferation, as assessed by Ki67 staining in insulin-positive cells, and reduced β-cell death, as assessed by TUNEL staining in insulin-positive cells (**Fig. 4H-I**). Together, these results indicate that H+E alone does not reverse recent-onset autoimmune diabetes but becomes effective when combined with transient anti-CD3-mediated immune modulation. This combined treatment improves glycemia, promotes diabetes remission, restores β-cell mass, increases β-cell proliferation, and reduces β-cell death in recent-onset diabetic NOD mice.

**Figure 4.**
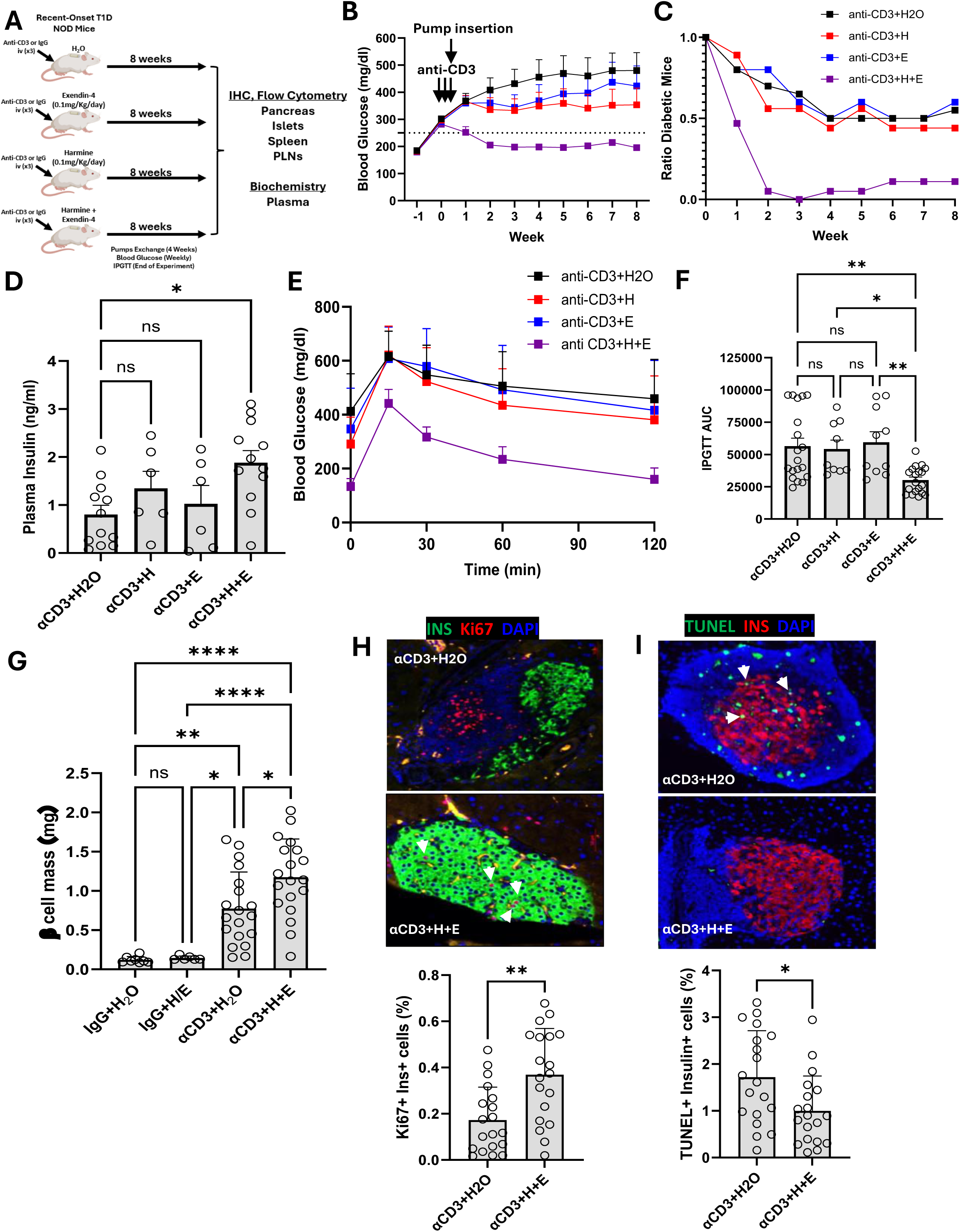
Treatment of recent-onset diabetic NOD mice with anti-CD3, harmine and exendin-4. **A** Scheme depicting the treatment of NOD mice. Created in BioRender. Garcia-Ocana, A. (2026). **B** Random blood glucose levels in recent onset NOD mice treated iv for 3 days with anti-CD3 (αCD3) and then implanted with Alzet minipumps to deliver vehicle (H_2_O), harmine (H), exendin-4 (E) or H+E for 56 days. Dash line indicates 250mg/dL blood glucose. N=19 αCD3+H_2_O-treated-mice, N=9 αCD3+H-treated mice, N=10 αCD3+E-treated mice, N=19 αCD3+H+E-treated mice. **C** Ratio of diabetic mice during the treatment period. **D** Non fasting plasma insulin levels at the end of treatment. One-way ANOVA with Tukey’s multiple comparisons test, *P<0.05. **E** Intraperitoneal glucose tolerance test and **F** area under the curve in these mice. One-way ANOVA with Tukey’s multiple comparisons test, *P<0.05 and **P<0.01. **G** β-cell mass analysis in pancreatic sections from IgG- or αCD3-pre-treated mice and then with H+E. One-way ANOVA with Tukey’s multiple comparisons test, *P<0.05, **P<0.01, ****P<0.0001. Representative images and quantitation (below) of β-cell **H** proliferation (Ki67) and **I** death (TUNEL) in αCD3-pre-treated mice and then with H_2_O or H+E. Two-tailed Student’s t-test, *P<0.05, **P<0.01.

### Harmine plus Exendin-4 reduces islet immune infiltration and increases local FoxP3+ regulatory T cells after anti-CD3 therapy

Since anti-CD3+H+E treatment improved glycemia and increased β-cell mass in recent-onset diabetic NOD mice, we next examined whether this response was associated with changes in islet immune infiltration. Histological analysis showed that anti-CD3+H+E treatment reduced the proportion of islets with severe insulitis and increased the proportion of islets with limited immune infiltration compared with control mice treated with anti-CD3 alone (**Fig. 5A**). To further characterize the local islet immune environment, we analyzed pancreatic sections from an independent 4-week treatment cohort of mice (**Suppl. Fig. 4A**). Pancreas immunostaining showed that anti-CD3+H+E markedly reduced CD45+ immune-cell infiltration surrounding insulin-positive islets (**Fig. 5B**). Density of B220+ B-cells showed a decreasing trend in anti-CD3+H+E-treated mice (**Fig. 5C**), while CD3+ T-cell density was significantly reduced (**Fig. 5D**). RNAscope analysis further showed decreased *Cd8a*+ and *Cd4*+ cells in islets from anti-CD3+H+E-treated mice compared with anti-CD3-treated controls (**Fig. 5E**). In contrast, *Foxp3+* regulatory T-cell signal relative to *Cd4+* cells was increased after anti-CD3+H+E treatment (**Fig. 5F**).

**Figure 5.**
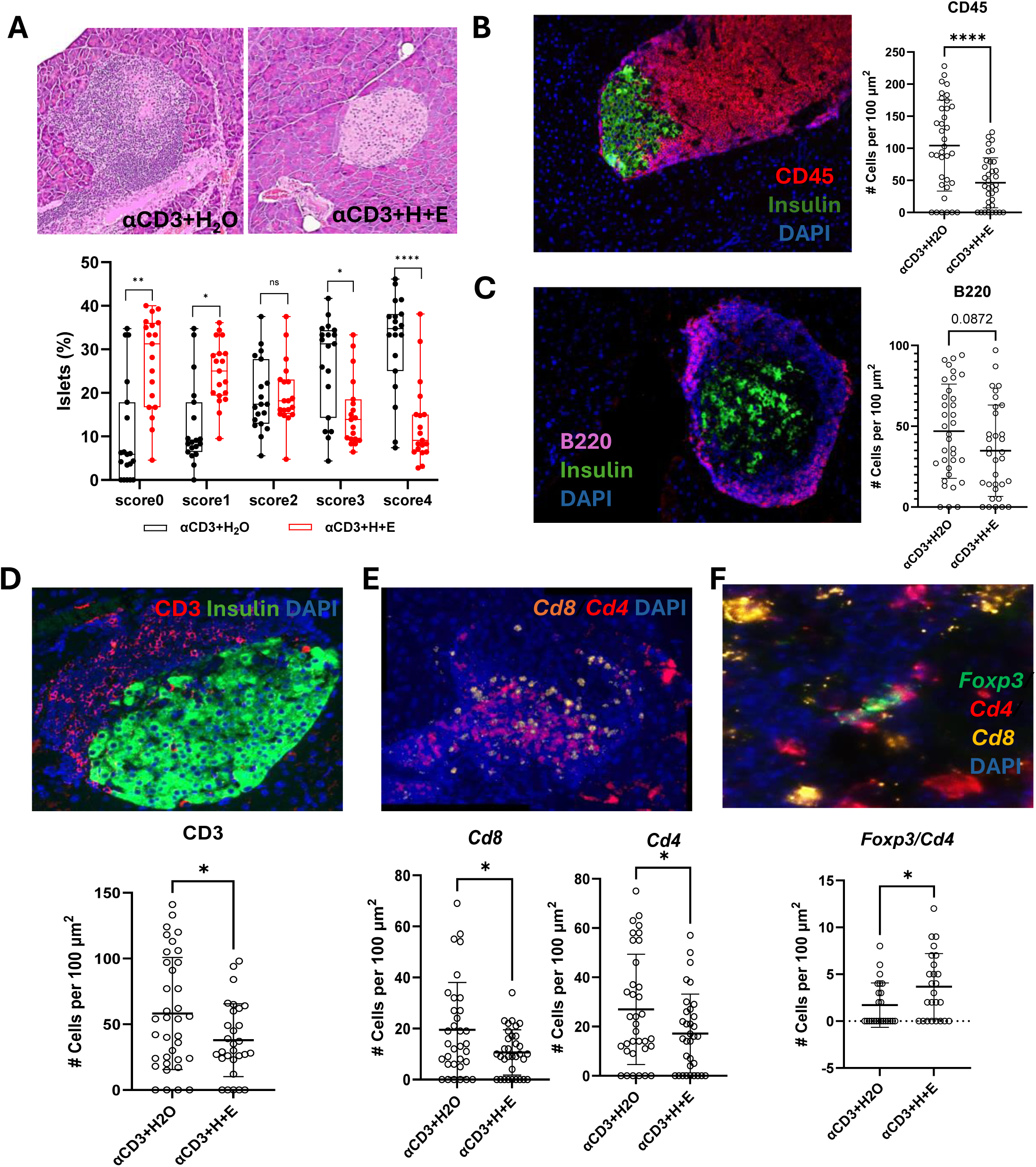
Insulitis in recent-onset diabetic NOD mice treated with anti-CD3, harmine and exendin-4. **A** Representative images and quantitation of insulitis in pancreatic sections from anti-CD3 (αCD3) and H_2_O or H+E treated mice, with scores ranging from 0 (no insulitis) to 4 (full insulitis). Two-tailed Student’s t-test, *P<0.05, **P<0.01, ****P<0.0001. Representative image of **B** CD45 (red) and insulin (green) immunolabeling, **C** B220 (magenta) and insulin (green) immunolabeling, and **D** CD3 (red) and insulin (green) of an islet from αCD3+H_2_O-treated mouse and quantitation of these cells per 100 µm2 area around insulin+ area in pancreatic sections from αCD3- and H_2_O or H+E treated mice. N=3 mice per group. Blue=DAPI. Two-tailed Student’s t-test, *P<0.05, ****P<0.0001. **E** Representative image of RNA Scope with probes for *Cd8* (orange), *Cd4* (red), **F** and *Foxp3* (green) of an αCD3+H_2_O-treated mouse and quantitation of *Cd8*+, *Cd4*+ and *Foxp3/Cd4*+ cells per 100 µm2 area around islet area in pancreatic sections from αCD3- and H_2_O or H+E treated mice. N=3 mice per group. Blue=DAPI. Two-tailed Student’s t-test, *P<0.05.

Consistent with the immunostaining observations, flow cytometry analysis of isolated islets from the 4-week cohort mice showed reduced percentage of CD45+ immune cells in anti-CD3+H+E-treated mice compared with anti-CD3-treated controls (**Suppl. Fig. 4A-C**). Together, these findings indicate that H+E treatment following low-dose anti-CD3 reduces local islet immune infiltration, decreases islet-associated CD4+ and CD8+ T-cell accumulation, and increases local Foxp3+ regulatory T-cell enrichment.

### Harmine plus Exendin-4 reduces T cell activation and increases regulatory T cells and T cell exhaustion markers in early-onset T1D NOD mice pre-treated with anti-CD3

Anti-CD3 therapy has shown promise in patients with recent-onset Type 1 Diabetes (T1D), in part by enhancing regulatory T cells (Tregs), reducing pathogenic T-cell responses, and promoting T-cell exhaustion-associated programs (10,11). Since low-dose anti-CD3 followed by H+E administration reversed recent-onset T1D in NOD mice (**Fig. 4**) and reduced CD4 and CD8 T-cell infiltration while increasing Treg presence in pancreatic islets (**Fig. 5**), we next examined whether H+E further modified peripheral lymphocyte composition and T-cell phenotypes in anti-CD3-treated mice. Anti-CD3+H+E treatment did not significantly alter the number of CD45+ cells or the CD4/CD8 T-cell ratios in the spleen or pancreatic lymph nodes (pLNs) compared with anti-CD3-treated mice at the end of treatment, day 56 (**Fig. 6A**, **Suppl. Fig. 5A-B**). The percentages of naïve, memory, and effector CD4 and CD8 T-cell populations, defined by CD44 and CD62L expression, were also largely unchanged in pLNs after anti-CD3+H+E treatment (**Fig. 6B**), except for a modest reduction in naïve CD4 T cells. These findings indicate that the improved response to anti-CD3+H+E was not associated with broad depletion of peripheral immune cells or major changes in the distribution of conventional T-cell subsets.

**Figure 6.**
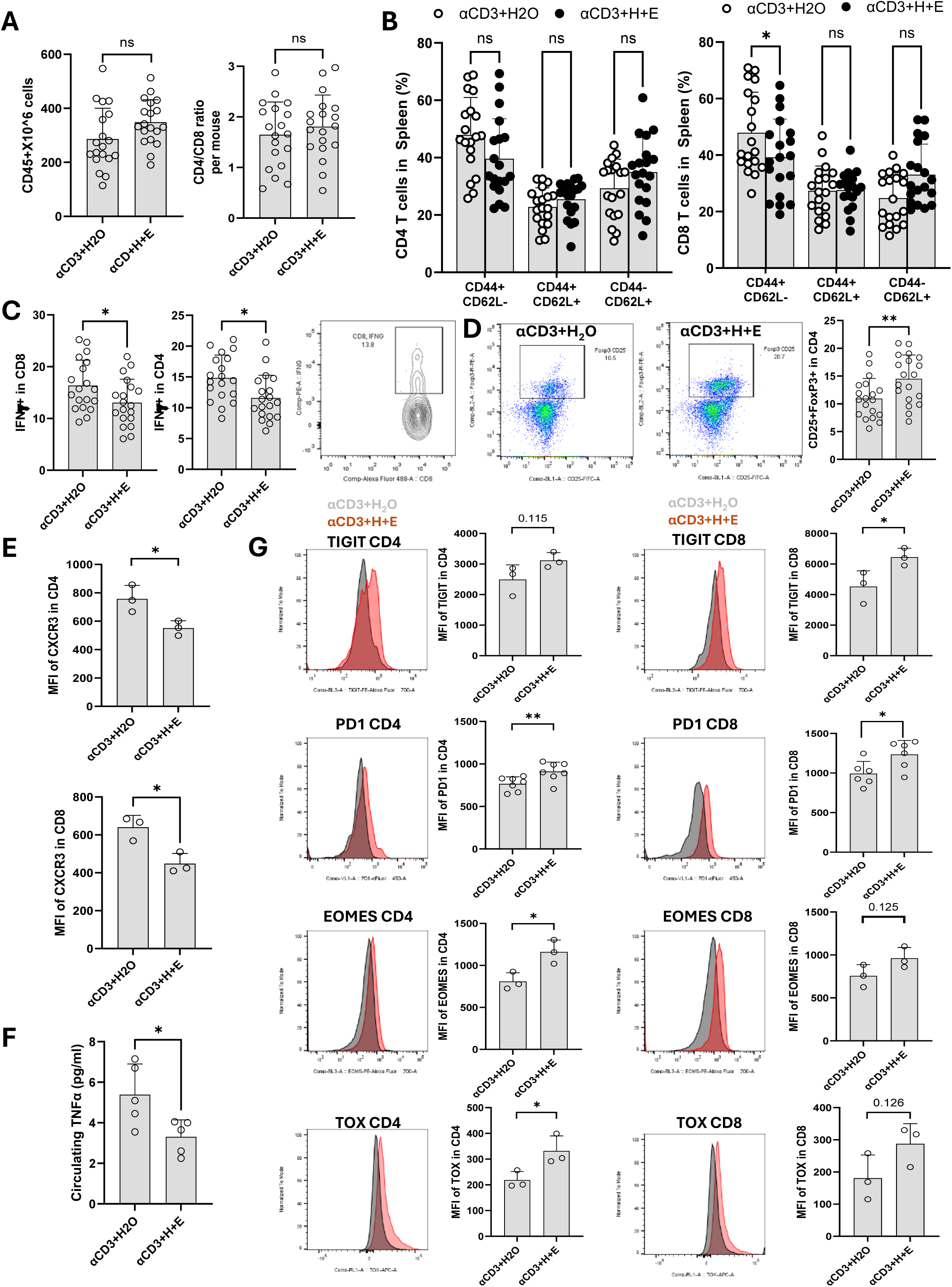
Immunophenotyping of recent-onset diabetic NOD mice treated with anti-CD3 and harmine and exendin-4. **A** Quantitation of CD45+ cells and the CD4/CD8 T-cell ratios in pancreatic lymph nodes (pLNs) in anti-CD3 (αCD3)-H_2_O- and αCD3-harmine (H)+Exendin-4 (E)-treated mice at the end of treatment (day 56). N=19/group. Two-tailed Student’s t-test. **B** Quantitation of naïve (CD44+CD62L-), memory (CD44+CD62L+), and effector (CD44-CD62L+) CD4 and CD8 T-cell populations in pLNs in αCD3-H_2_O- and αCD3-H+E-treated mice at the end of treatment (day 56). N=19/group. Two-tailed Student’s t-test, *P<0.05. **C** Quantitation of IFN-γ-producing CD4+ and CD8+ T cells in pLNs after ex vivo stimulation in αCD3-H2O- and αCD3-H+E-treated mice at the end of treatment (day 56). N=19/group. Two-tailed Student’s t-test, *P<0.05. **D** Representative flow cytometry plots and quantitation of CD25+FoxP3+ CD4 T cells in αCD3-H2O- and αCD3-H+E-treated mice at the end of treatment (day 56). N=19/group. Two-tailed Student’s t-test, *P<0.05. **E** Quantitation of CXCR3 expression in both CD4 and CD8 T cells in a subset of mice treated 4 weeks with αCD3-H2O or αCD3-H+E. N=3. Two-tailed Student’s t-test, *P<0.05. **F** Circulating levels in αCD3-H2O-and αCD3+H+E-treated mice at the end of the 4-week treatment. N=3. Two-tailed Student’s t-test, *P<0.05. **G** Quantitation of T-cell exhaustion-associated markers TIGIT, PD1, EOMES, and TOX in splenic CD4 and CD8 T cells from mice treated for 28 days with αCD3-H2O or αCD3+H+E. N=3-7 mice per group. Two-tailed Student’s t-test, *P<0.05.

We next assessed whether the remaining T cells displayed altered inflammatory function. After *ex vivo* stimulation, the frequencies of IFN-γ-producing CD4+ and CD8+ T cells in pLNs were significantly decreased in anti-CD3+H+E-treated NOD mice compared with anti-CD3-treated controls (**Fig. 6C**). In parallel, the percentage of CD25+FoxP3+ CD4 T cells was significantly increased in anti-CD3+H+E-treated mice (**Fig. 6D**). Thus, addition of H+E to anti-CD3 reduced inflammatory T-cell responses while increasing regulatory T-cell frequency. We then examined CXCR3, a chemokine receptor involved in T-cell migration during inflammatory immune responses. Mice treated for 4 weeks with anti-CD3+H+E (**Suppl. Fig. 4A**) displayed significantly decreased CXCR3 expression in both CD4 and CD8 T cells in the spleen compared with anti-CD3-treated mice (**Fig. 6E**). Circulating levels of TNFα were also significantly reduced in anti-CD3+H+E-treated mice compared with anti-CD3-treated controls at the end of the 4-week treatment (**Fig. 6F**).

Next, we analyzed T-cell exhaustion-associated markers in splenic CD4 and CD8 T cells from mice treated for 28 days with anti-CD3+H+E or anti-CD3 alone. Expression of TIGIT, PD1, EOMES, and TOX was increased in CD4 and CD8 T cells from anti-CD3+H+E-treated mice compared with anti-CD3-treated controls (**Fig. 6G**). Taken together, these results indicate that H+E treatment following low-dose anti-CD3 does not induce broad peripheral lymphocyte depletion but reduces IFN-γ-producing T cells, increases Tregs, decreases CXCR3 expression, reduces circulating TNFα, and enhances exhaustion-associated T-cell marker expression. These immune changes are consistent with reduced islet infiltration and improved diabetes remission in anti-CD3+H+E-treated NOD mice.

### Harmine plus Exendin-4 diminishes T cell activation, increases Tregs and enhances T cell exhaustion markers in human PBMCs

The immune changes observed in anti-CD3+H+E-treated NOD mice could result from direct effects of H+E on immune cells, indirect effects secondary to improved β-cell survival and reduced islet inflammation, or systemic changes associated with improved glycemia. To determine whether H+E could directly modulate human immune cells, we performed Cytometry by Time-of-Flight (CyTOF) analysis of human peripheral blood mononuclear cells (PBMCs) treated *in vitro*. PBMCs from three healthy donors were divided into four aliquots and treated for 6 days with vehicle, H, E, or H+E in the presence of low-dose anti-CD3/anti-CD28 and IL2/IL7 stimulation (**Fig. 7A**). CyTOF data were analyzed using both algorithm-based analysis and conventional FlowJo analysis. After embedding the CyTOF data into Seurat, we performed integration and visualization and confirmed comparable distribution of cells across treatment groups (**Suppl. Fig. 6A-F**). Major immune-cell populations were assigned using established markers, including CD8+ T cells, CD4+ T cells, NK cells, B cells, and myeloid cells (**Suppl. Fig. 6C-F**). We first analyzed activation-associated markers in T-cell populations. In CD4+ non-Treg cells, H and H+E reduced expression of CD69, CD127, and CD27 and decreased the combined activation score (**Fig. 7B**). A similar reduction in activation-associated markers and combined activation score was observed in CD8+ T cells treated with H or H+E (**Fig. 7B**). These data indicate that harmine-containing treatments suppress activation-associated phenotypes in stimulated human T cells.

**Figure 7.**
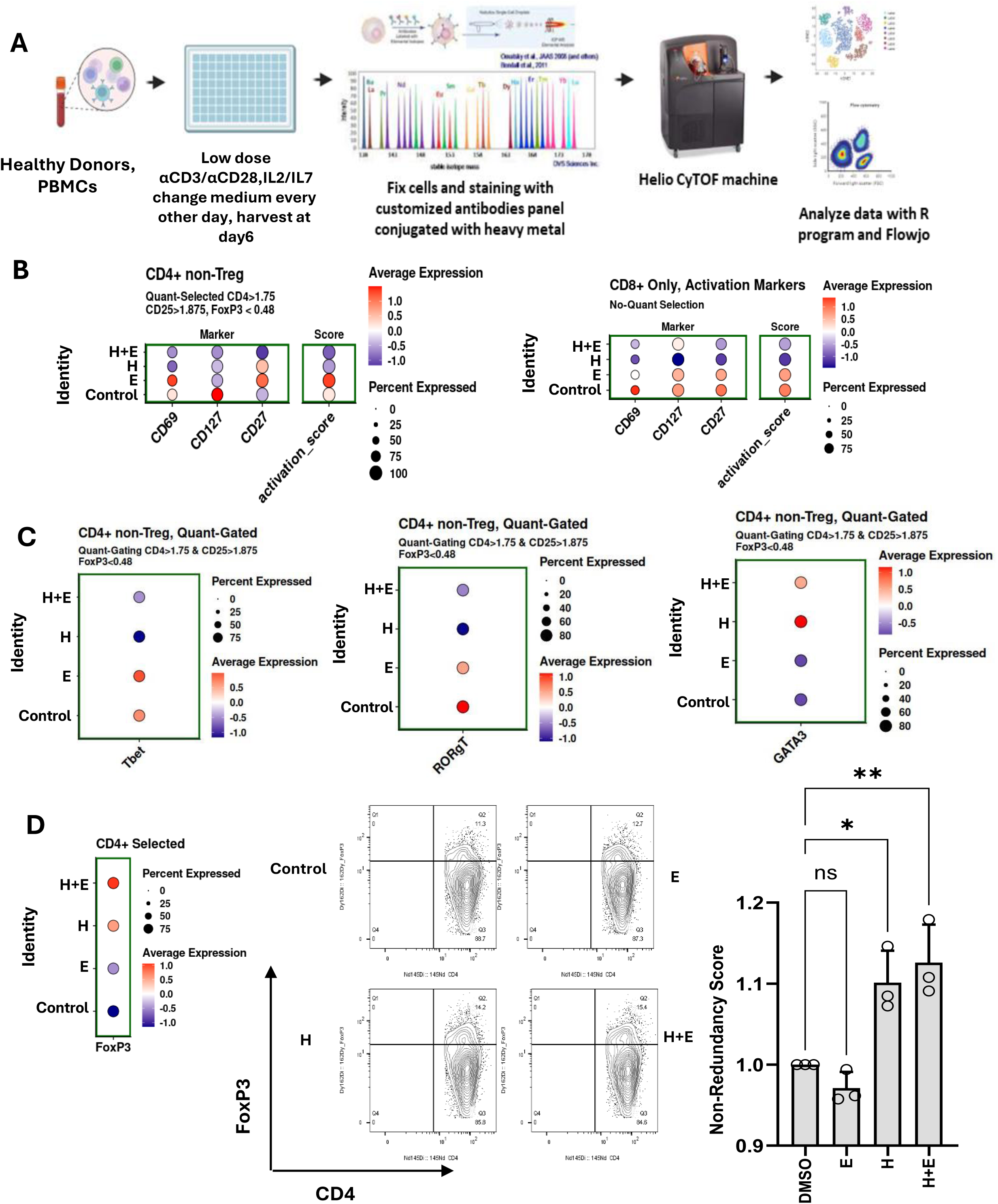
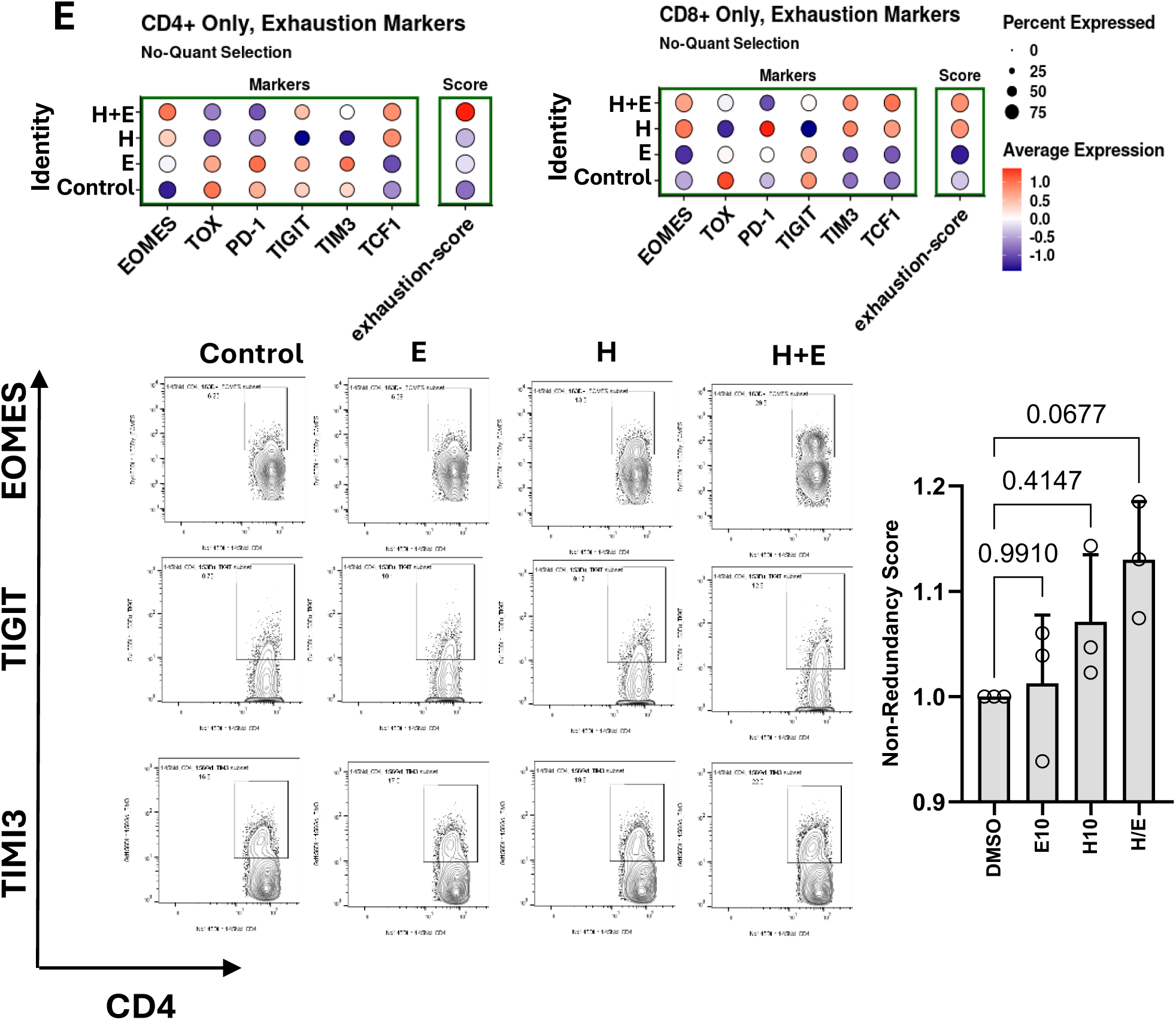
CyTOF analysis of human PBMCs treated with harmine and exendin-4. **A** Schematic representation of the experiments. Created in BioRender. Garcia-Ocana, A. (2026). After embedding the CyTOF data into Seurat, we performed integration, visualization and expression analysis. **B** Dot plot represents expression of CD69, CD127 and CD27 in non-Treg CD4+ cells (left) and CD8+ cells (right) in the four treatment groups: harmine (H), Exendin-4 (E) and H+E. Activation scores of these immune cells based on the indicated markers appear next to it. **C** Dot plots represent expression of TBET (Th1), RORgT (Th17) and GATA3 (Th2) in non-Treg CD4+ cells in the four treatment groups. **D** Dot plot represents expression of FOXP3 in CD4+ cells (left) and FlowJo-based analysis of the CyTOF data in the four treatment groups confirming increases in Treg cells in H and H+E groups. One-way ANOVA with Tukey’s multiple comparisons test, *P<0.05, **P<0.01. **E** Dot plots represent expression of T cell exhaustion markers EOMES, TOX, PD-1, TIGIT, TIM3, and TCF1 CD4+ cells (left) and CD8+ cells (right) in the four treatment groups. Activation score of these immune cells based on the indicated markers appear next to it. FlowJo-based analysis of the CyTOF data (below) for representative exhaustion markers such as EOMES, TIGIT and TIM3 in the four treatment groups confirming increases in exhaustion markers by H+E. One-way ANOVA with Tukey’s multiple comparisons test.

We next examined helper T-cell lineage-associated markers in CD4+ non-Treg cells. H+E reduced Tbet and RORγT expression, suggesting decreased Th1 and Th17-associated polarization, while GATA3 expression was increased, suggesting a relative shift toward a Th2-associated phenotype (**Fig. 7C**). Analysis of FoxP3 expression showed increased CD4+FoxP3+ Tregs in H- and H+E-treated PBMCs, but not in E-treated cells (**Fig. 7D**). This finding was confirmed by FlowJo-based analysis of the CyTOF data (**Fig. 7D**). We then assessed exhaustion-associated markers in CD4+ and CD8+ T cells. In CD4+ T cells, H+E increased several exhaustion-associated markers, including EOMES, TIGIT, TIM3, TOX, TCF1, and PD1, resulting in an increased exhaustion score (**Fig. 7E**). A similar increase in exhaustion-associated marker expression and exhaustion score was observed in CD8+ T cells (**Fig. 7E**). FlowJo-based analysis further confirmed increased expression of selected exhaustion-associated markers, including EOMES, TIGIT, and TIM3, in H- and H+E-treated cells (**Fig. 7E**). Together, these results indicate that H+E directly modulates activated human PBMCs by reducing T-cell activation, decreasing Th17-associated features, increasing FoxP3+ Tregs, and enhancing exhaustion-associated/inhibitory T-cell phenotypes. These findings are consistent with the immune phenotype observed in anti-CD3+H+E-treated NOD mice and support the possibility that H+E contributes to immune modulation in addition to its β-cell protective effects.

### SNHG6 is an H+E-induced β-cell lncRNA that contributes to β-cell protection under conditions of inflammatory stress

The preceding studies showed that H+E suppresses inflammatory, immunogenicity, and stress-related programs in cytokine-treated human β-cells. We next sought to identify β-cell-intrinsic genes selectively regulated by H+E under inflammatory conditions. Using the human islet scRNA-seq dataset in **Fig. 1**, we first identified 288 differentially expressed genes enriched in β-cells treated with H+E in the presence of cytokines compared with cytokine-treated controls (**Supplementary Data 1**). Genes shared with H- or E-treatment conditions, as well as genes enriched in non-β cell populations, were then excluded to identify H+E-selective β-cell genes. This filtering strategy identified 64 β-cell-selective H+E-regulated genes (**Supplementary Data 1**), of which 8 were expressed in more than 40% of β-cells under control conditions: *SNHG6, TMA7, MYL6B, PPP1R14B, MNX1, TMEM14A, ZKSCAN1,* and *MRPS34* (**Fig. 8A-B**).

**Figure 8.**
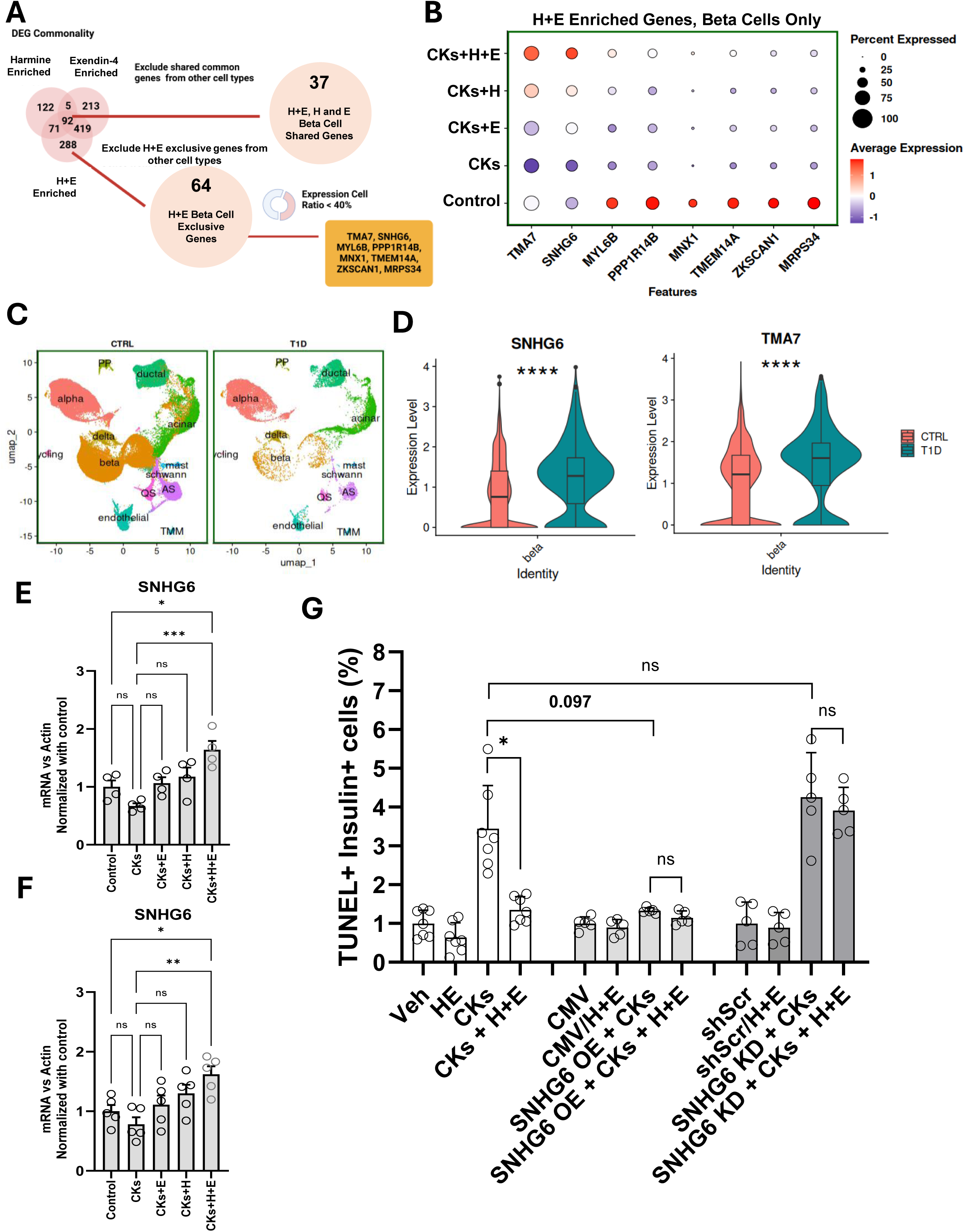
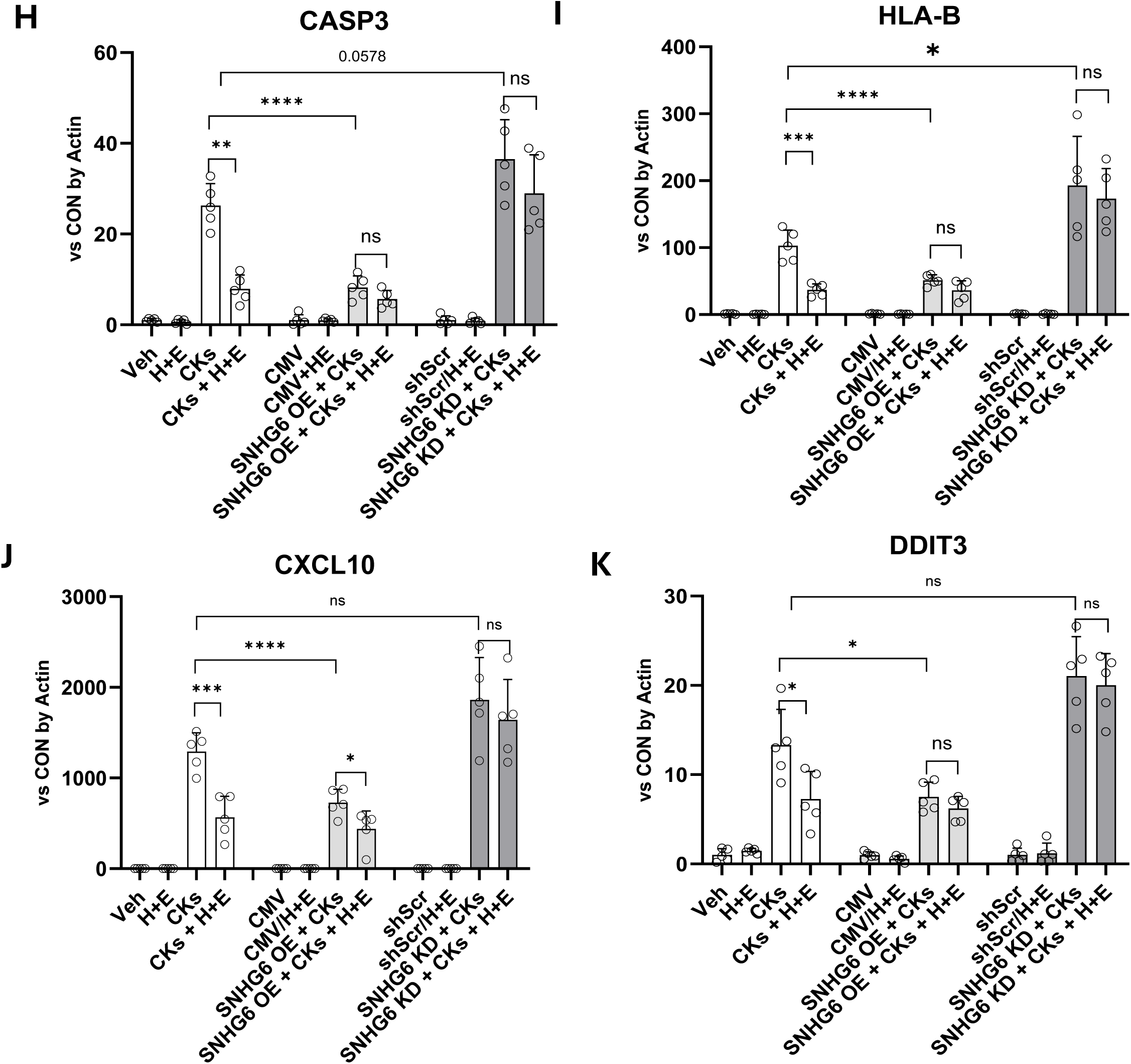
scRNA-seq analysis of human islets under cytokines, harmine and exendin-4 treatment uncovers SNHG6 as a mediator of β-cell survival and immunogenicity. **A.** Gene commonality analysis of differentially expressed genes (DEGs) in harmine (H), exendin-4 (E) and H+E vs vehicle in cytokine (Cks)-treated human islets. Of the 92 common genes, 37 genes are common in β-cells. Of the 288 DEGs only in H+E-treated islet cells, 64 genes are excusive for β-cells. Of these, only 8 are expressed in more than 40% of the β-cells. **B** Dot plot represents gene expression of the 8 commonality genes in the different treatment groups. **C** Unsupervised clustering, cell type annotated UMAP of scRNA-seq data set from HPAP-HIRN of human islets from non-diabetic and T1D human donors. **D** Violin plots represent the expression of *SNHG6* and *TMA7* in β-cells in the HPAP-NIRN dataset. Wilcoxon rank-sum test, P<0.0001. **E** SNHG6 expression in human islets treated with cytokines and vehicle, H, E and H+E compared with non-cytokine treated cells assessed by qPCR. N=4 different human islet preparations. One-way ANOVA with Tukey’s multiple comparisons test, *P<0.05, ***P<0.001. **F** SNHG6 expression in EndoC-βH1 human β-cells treated with cytokines and vehicle, H, E and H+E compared with non-cytokine treated cells assessed by qPCR. N=5 experiments. One-way ANOVA with Tukey’s multiple comparisons test, *P<0.05, **P<0.01.**G** Quantitation of human β-cell death in EndoC-βH1 cells assessed by TUNEL and insulin immunostaining. EndoC-βH1 cells were treated with vehicle, H+E, cytokines and cytokines+H+E in non-transduced cells or transduced with Adv-CMV (control), Adv-SNHG6, Adv-scramble shRNA, or Adv-shSNHG6 (SNHG6 knockdown, KD). N=5-7 experiments. One-way ANOVA with Tukey’s multiple comparisons test, *P<0.05. mRNA expression of **H** *CASP3*, **I** *HLA-B*, **J** *CXCL10*, and **K** *DDIT3* (CHOP) in EndoC-βH1 human β-cells treated as in G. N=5 experiments. One-way ANOVA with Tukey’s multiple comparisons test, *P<0.05, **P<0.01, ***P<0.001, ****P<0.0001.

Among these candidates, *TMA7* and *SNHG6* showed the strongest induction in H+E-treated β-cells. To determine whether these candidates were associated with β-cells exposed to autoimmune inflammatory stress *in vivo*, we analyzed the human T1D scRNA-seq dataset from HIRN-HPAP (32). Both *SNHG6* and *TMA7* were increased in β-cells from T1D donors compared with control β-cells (**Fig. 8C-D**). We focused subsequent studies on SNHG6 because it is a long non-coding RNA with reported roles in cellular stress and survival pathways, whereas its function in human β-cells remains unknown. Validation of SNHG6 expression regulation by qPCR showed that H+E increased SNHG6 expression compared with cytokine-treated controls and individual drug treatments in human islets (**Fig. 8E**) and EndoC-βH1 cells (**Fig. 8F**). These findings identify SNHG6 as an H+E-induced β-cell lncRNA associated with inflammatory stress conditions.

Having identified SNHG6 as an H+E-regulated β-cell lncRNA, we next tested whether SNHG6 functionally regulates β-cell responses to inflammatory cytokines. EndoC-βH1 cells were transduced to overexpress SNHG6 or to reduce SNHG6 expression (**Suppl. Fig. 7**), followed by cytokine treatment in the presence or absence of H+E. SNHG6 overexpression reduced cytokine-induced β-cell death (**Fig. 8G**), and *CASP3* expression (**Fig. 8H**) and decreased expression of *HLA-B* (**Fig. 8I**) and *CXCL10* (**Fig. 8J**), two immunogenicity/inflammatory-related genes suppressed by H+E in cytokine-treated human β-cells. SNHG6 overexpression also reduced *DDIT3* (CHOP) expression, suggesting attenuation of cytokine-induced stress-response signaling (**Fig. 8K**). Conversely, SNHG6 knockdown increased cytokine-induced *CASP3, HLA-B, CXCL10*, and *DDIT3* (CHOP) expression and reduced the suppressive effect of H+E on these cytokine-induced markers and β-cell death (**Fig. 8G-K**). Together, these gain- and loss-of-function studies indicate that SNHG6 contributes to the ability of H+E to suppress β-cell apoptotic, stress-related, and immunogenicity-associated responses under inflammatory conditions.

## Discussion

In this study, we demonstrate that a β-cell regenerative and protective therapy combining harmine and exendin-4 (H+E) is highly effective in reversing recent-onset autoimmune diabetes when paired with transient immune modulation using anti-CD3. In NOD mice with recent-onset diabetes, treatment with anti-CD3 followed by H+E improved glycemic control, increased the rate of diabetes remission, enhanced plasma insulin levels and glucose tolerance, promoted β-cell proliferation and β-cell mass expansion, and reduced β-cell death. These findings support a model in which anti-CD3 establishes a permissive immunological environment that enables H+E to drive functional β-cell recovery (6–11). Collectively, our results demonstrate the synergistic potential of combining immune modulation with β-cell regenerative therapies and, to our knowledge, represent the first evidence that this strategy can induce durable remission of recent-onset autoimmune diabetes.

This finding reinforces the concept that immune modulation alone may be insufficient to achieve durable remission of T1D, particularly once substantial β-cell loss has already occurred. Although anti-CD3 therapies such as teplizumab can preserve residual β-cell function and delay disease progression (6–9), their clinical benefit has generally been transient, likely reflecting the limited capacity of the remaining β-cell mass to restore normoglycemia in the setting of established disease. Increasing evidence indicates that anti-CD3-mediated efficacy extends beyond transient immunosuppression and involves broader reprogramming of the autoimmune response, including expansion of regulatory T-cell populations, attenuation of pathogenic effector T-cell activity, and induction of exhaustion-associated transcriptional programs in autoreactive CD8+ T cells that correlate with improved clinical outcomes (10,11). In the present studies, we employed a low-dose anti-CD3 regimen (5 μg/mouse for 3 days) that has previously been shown to have limited efficacy in inducing remission of recent-onset diabetes in NOD mice (31). Our results suggest that combining low-dose anti-CD3 with the β-cell regenerative and protective regimen of harmine plus exendin-4 overcomes key limitations of anti-CD3 monotherapy by simultaneously targeting both the immune and β-cell compartments of the disease. Beyond increasing β-cell mass, H+E may reduce β-cell immunogenicity and antigen-processing pathways, thereby diminishing the antigenic stimuli that sustain autoimmune responses. This concept is consistent with expanding evidence that β cells actively participate in T1D pathogenesis through enhanced antigen presentation, MHC upregulation, and interactions with autoreactive T cells, rather than serving merely as passive targets of immune attack (15–19). In this context, transient immune intervention may create a therapeutic window during which newly generated and surviving β cells are protected from immune-mediated destruction, allowing regeneration and functional recovery to proceed. The marked reversal of recent-onset autoimmune diabetes observed with the combined treatment supports a model in which successful disease modification requires coordinated restoration of immune tolerance and β-cell mass. These findings further suggest that anti-CD3-based therapies may achieve greater and more durable efficacy when incorporated into combination approaches that actively promote β-cell regeneration, rather than relying solely on preservation of the limited endogenous β-cell reserve. Such strategies could represent an important step toward achieving sustained remission and, potentially, functional cure in individuals with recent-onset T1D.

The immune effects observed in anti-CD3+H+E-treated NOD mice were not attributable to broad peripheral lymphocyte depletion. In these studies, we used a low dose of anti-CD3 (31) that largely preserved total CD45+ cell numbers, CD4/CD8 T-cell ratios, and major naïve, memory, and effector T-cell populations. Instead, anti-CD3+H+E selectively attenuated inflammatory T-cell activity, as evidenced by reduced frequencies of IFN-γ-producing CD4+ and CD8+ T cells, expansion of CD25+FoxP3+ regulatory T cells, decreased CXCR3 expression, lower circulating TNFα levels, and increased expression of exhaustion-associated T-cell markers. These findings suggest that H+E enhances anti-CD3-mediated remodeling of autoreactive immune responses (10,11) by shifting T cells away from a highly inflammatory, diabetogenic phenotype toward a more regulated and functionally restrained state. Consistent with these observations, similar effects were observed in activated human PBMCs treated *in vitro*, where H+E reduced T-cell activation, increased the frequency of FoxP3+ regulatory T cells, suppressed Th17-associated features, and upregulated exhaustion-associated and inhibitory markers. Collectively, these human immune cell data support the possibility that anti-CD3+H+E can directly modulate immune-cell phenotypes. However, studies using PBMCs from individuals with T1D will be required to determine whether these immunomodulatory effects are maintained in disease-relevant human immune cells.

In addition to its immunomodulatory effects and the expected enhancement of β-cell proliferation (12–14,29,33–35), results indicate that H+E also increases β-cell resilience against autoimmune-mediated injury. Previous studies have shown that H+E improves β-cell survival following islet transplantation (14). Extending these observations, we find that H+E suppressed the expression of inflammatory and pro-apoptotic genes, reduced the activation of cytokine-responsive IFNγ-, IL1/IL1R-, and TNFα-associated pathways, and decreased β-cell death in cytokine-treated human islets. H+E also reduced *iNOS* expression, leading to lower NO production, a key mediator of β-cell dysfunction and death (36,37). Together, these findings suggest that H+E not only promotes β-cell proliferation and survival under non-inflammatory conditions but also directly mitigates inflammatory signaling pathways that contribute to β-cell injury in T1D.

A second important β-cell effect of H+E was the suppression of cytokine-induced immunogenicity programs. Exposure of human β-cells to inflammatory cytokines induced the expression of classical MHC class I genes, IFN-responsive transcription factors, antigen-processing and presentation machinery, and chemokines, consistent with previous reports demonstrating that inflammatory stress drives β-cell acquisition of a more immunogenic phenotype in T1D (17–19). Specifically, cytokine treatment increased the expression of *HLA-A, HLA-B, HLA-C, B2M*, IFN-regulated transcription factors of the IRF family, antigen-processing genes including *PSMB8, PSMB9, PSMB10*, and *TAP1*, and the T-cell chemoattractants *CXCL9, CXCL10, and CXCL11*. H+E markedly attenuated these responses while increasing expression of the non-classical MHC class I molecule HLA-E, which has been implicated in immune regulation and protection from immune-mediated cytotoxicity (38,39). These findings suggest that H+E may reduce β-cell immune visibility while simultaneously limiting the production of signals that promote immune-cell recruitment under inflammatory conditions. Pro-inflammatory cytokines can enhance antigen presentation, increase MHC class I expression, and induce chemokine production, thereby amplifying immune-cell infiltration and local inflammation within islets (17–19). The concurrent reduction of *CXCR3* expression on T cells *in vivo* and *CXCL9/CXCL10/CXCL11* expression in cytokine-treated β-cells suggests that anti-CD3+H+E may disrupt the CXCR3-chemokine axis that contributes to T-cell trafficking into inflamed islets (40,41). By simultaneously targeting immune-cell migratory capacity and β-cell-derived chemotactic signals, anti-CD3+H+E may interrupt a feed-forward cycle of islet inflammation, immune-cell recruitment, and progressive β-cell destruction that characterizes T1D pathogenesis (17–19,38–41).

H+E also improved β-cell stress responses and secretory function under inflammatory conditions. Exposure of human β-cells to pro-inflammatory cytokines activated cellular stress pathways, including unfolded protein response (UPR) and endoplasmic reticulum (ER) stress programs, which are well-established contributors to β-cell dysfunction and death in T1D (42,43). H+E significantly reduced the activity of these stress-related pathways. qPCR analysis confirmed decreased expression of *ATF6* and spliced *XBP1*, key mediators of canonical UPR signaling, as well as *ATF3*, *TXNIP*, and *DDIT3* (CHOP), markers associated with integrated stress responses, inflammation-induced β-cell injury, oxidative stress, and maladaptive ER stress leading to apoptosis (42,43). These findings indicate that H+E attenuates multiple convergent stress pathways that contribute to β-cell failure during autoimmune inflammation. Importantly, the reduction in cellular stress was accompanied by partial preservation of genes essential for β-cell identity and function, including *INS*, *CHGA*, *PCSK1*, and *GLP1R* (44). H+E treatment also improved insulin secretory quality, as reflected by reduced proinsulin release and a lower proinsulin-to-insulin ratio. This observation is particularly noteworthy because impaired proinsulin processing and elevated circulating proinsulin levels are recognized features of stressed and dysfunctional β-cells in individuals with T1D and in prediabetic states (45,46). Together, these findings suggest that H+E not only enhances β-cell survival under inflammatory conditions but also preserves functional β-cell competence by reducing stress-induced defects in hormone processing and secretion. By simultaneously limiting inflammatory injury, ER stress, and secretory dysfunction, H+E may help maintain a healthier and more resilient β-cell phenotype during autoimmune attack.

Our single-cell transcriptomic analysis further identified SNHG6 as an H+E-induced β-cell lncRNA under inflammatory conditions. *SNHG6* expression in human β-cells was suppressed by cytokine treatment and upregulated by H+E. Its upregulation was confirmed in cytokine-treated human islets and EndoC-βH1 cells, and increased expression was also observed in β-cells from donors with T1D. SNHG6 is a highly conserved lncRNA that has been implicated in the regulation of cellular stress responses, survival pathways, proliferation, apoptosis, and inflammatory signaling in multiple tissues and disease contexts, including metabolic disorders and diabetes-related complications (25,26,47,48). Mechanistically, SNHG6 has been reported to function as a competing endogenous RNA, modulating gene-expression networks through interactions with microRNAs and downstream stress-response pathways (25,26,47–49). Although its role in pancreatic β-cells remains largely unexplored, the observed cytokine-mediated suppression and H+E-induced restoration of SNHG6 expression suggest that this lncRNA may participate in adaptive responses that promote β-cell survival and preserve cellular homeostasis during inflammatory stress. Indeed, functional studies showed that SNHG6 overexpression reduced cytokine-induced *CASP3, HLA-B, CXCL10*, and *DDIT3* (CHOP) expression, while SNHG6 knockdown increased these inflammatory, immunogenicity, and stress-associated markers and reduced the suppressive effect of H+E. These findings identify SNHG6 as a potential mediator of the β-cell protective effects of H+E and warrant future studies to define its functional role in β-cell stress adaptation, immune signaling, and survival in T1D.

Several limitations of the current study should be acknowledged. First, although NOD mice represent a well-established model of spontaneous autoimmune diabetes (50,51), they do not fully recapitulate the complexity and heterogeneity of human T1D, warranting caution when extrapolating these findings to the clinical setting. Second, the durability of anti-CD3+H+E-induced T1D remission remains unknown and will require longer-term follow-up studies. Third, the human PBMC experiments were conducted using cells from healthy donors under *in vitro* activation conditions. Future studies using PBMCs, autoreactive T cells, or other immune-cell populations derived from individuals with T1D will be important to determine the translational relevance of these findings. Fourth, while cytokine-treated human islets provide a useful model of inflammatory β-cell stress (18,19), they do not fully capture the cellular interactions and microenvironmental complexity of autoimmune islets *in vivo*, which limits interpretation of the observed effects on β-cell biology in the T1D context. Fifth, harmine alone exhibited beneficial effects across several of the *in vitro* parameters evaluated, suggesting that higher doses or the development of more potent harmine-derived small molecules (52), either as monotherapy or in combination with αCD3, may ultimately be capable of inducing remission in T1D. Finally, although SNHG6 gain- and loss-of-function studies support a functional role for this lncRNA in β-cell protection, additional mechanistic studies are needed to establish whether SNHG6 is required for the full protective effects of H+E in primary human β-cells and *in vivo* models of autoimmune diabetes. Despite these limitations, the complementary findings across NOD mice, human immune cells, and human islets provide convergent evidence supporting the therapeutic potential of anti-CD3+H+E as a strategy to simultaneously target autoimmune inflammation and β-cell vulnerability in T1D.

Together, these findings support a dual-system therapeutic model for recent-onset T1D. In this model, low-dose anti-CD3 reduces autoimmune pressure and establishes a more permissive immune environment by dampening pathogenic T-cell responses and promoting immune regulation, while H+E simultaneously enhances β-cell recovery and resilience. Specifically, H+E suppresses inflammatory and apoptotic signaling, reduces β-cell immunogenicity, improves cellular stress responses and secretory quality, and induces protective β-cell-intrinsic programs, including the lncRNA SNHG6. By targeting both the immune system and the β-cell compartment, anti-CD3+H+E addresses two fundamental drivers of disease progression: persistent autoimmunity and β-cell vulnerability. This integrated approach may provide a framework for combining transient immune modulation with β-cell regenerative and protective therapies, thereby promoting more durable remission and improved preservation of endogenous β-cell function in autoimmune diabetes.

## Acknowledgements

We thank the NIDDK-funded Integrated Islet Distribution Program (IIDP) at City of Hope and Prodo Labs for supplying human cadaveric islets. We thank Dr. Brian Armstrong at City of Hope Light Microscopy and Digital Imaging Core for helping with fluorescent microscopy. We acknowledge support from the Arthur-Riggs Diabetes and Metabolism Research Institute (ARDMRI) at City of Hope, the Wanek Family Fund, NIH grants R01 DK141874, R01 DK126450, R01 DK105015, U24DK098085 and a Pilot and Feasibility Grant (GL) from the Einstein-Sinai DRC (P30 DK020541). We thank the Human Pancreas Analysis Program (HPAP) Database (RRID:SCR_016202), consortia under Human Islet Research Network (RRID:SCR_014393) (https://hpap.pmacs.upenn.edu/) funded by the NIH grants UC4-DK112217 and UC4-DK112232 for providing scRNA-seq data from human islets from non-diabetic and T1D donors.

## Declaration of Interests

G.L., P.W., R.J.D, A.F.S. and A.G.-O. are inventors on patents filed by The Icahn School of Medicine at Mount Sinai. PaulexBio has licensed this patent portfolio from The Icahn School of Medicine at Mount Sinai. R.J.D, A.F.S. and A.G.-O. are members of the Scientific Advisory Board of Paulex Bio.

## Materials and Methods

### Human islet samples

Adult human pancreata and pancreatic islets from brain-dead donors were provided by Prodo Laboratories (Aliso Viejo, CA) according to the standard procedure and used for the studies described here (**Supplementary Table 1**). Briefly, islets were harvested from pancreata from deceased organ donors without any identifying information and with informed consent properly and legally secured, and Western Institutional Review Board (WIRB) approval. In addition, we mined the raw FASTQ data from the Human Pancreas Analysis Program (HPAP) database, consortia under the Human Islet Research Network (HIRN) and performed analysis of 28 non-diabetic and 9 T1D islet scRNA-sequencing datasets through secure file transfer protocol (SFTP) (details below) (32).

### Human islet cell processing following treatments

Human islets from three different cadaveric donors (3000 IEQs/donor) were used in these studies (**Supplementary Table 1**). Briefly, human islets were treated for 6h in RPMI medium in the presence or absence of recombinant human cytokines: IL1β (500 units/mL), IFNγ (1,000 units/mL), and TNFα (1,000 units/mL) and with vehicle (0.1% DMSO), H (10µM), E (10nM) or with H+E. Islet cells were then dissociated using pre-warmed Accutase (cat# 25–058-CL, Corning) and resuspended in binding buffer (cat# 130–090-101, Miltenyi Biotec) with dead cell removal beads and applied onto the dead cell removal column (cat # 130–042-401, Miltenyi Biotec), which was attached to the MACS separator. Then the cell concentration was measured with the Countess-3 Automated Cell Counter (Thermo-Fisher).

### Single-cell RNA sequencing (scRNA-seq), alignment, and matrix generation

Cells were prepared according to the 10x Genomics Single Cell 3′ Gene Expression v3.1 protocol, partitioned and barcoded using a 10x Genomics Chromium Controller, and processed for cDNA and library generation. Libraries were sequenced on an Illumina NovaSeq 6000. Raw sequencing reads were aligned and quantified with the 10x Genomics Cell Ranger count pipeline against the GRCh38-2020-A human reference. Fourteen libraries were processed with Cell Ranger v7.0.1, whereas HP21251_H10CK was processed with Cell Ranger v7.1.0. Ambient RNA was removed from each raw feature-barcode matrix using the CellBender remove-background command. For each library, the Cell Ranger Estimated Number of Cells was used as the initial value for --expected-cells and checked against the knee of the rank-ordered UMI plot. The --total-droplets-included value was selected to include all barcodes that could contain cells and extend into the empty-droplet plateau. CellBender was run with a false-positive rate of 0.01 for 150 epochs using GPU acceleration. The resulting background-corrected HDF5 matrices were used for downstream analysis.

### scRNA-seq quality control, integration, and pathway analysis

Background-corrected HDF5 matrices were imported into R using Read10X_h5 and converted to Seurat objects. Genes expressed in fewer than three cells and cells with fewer than 200 detected genes were excluded during object creation. Sample objects were merged, and library complexity was calculated as log10(detected genes)/log10(UMIs). Cells with fewer than 500 UMIs, fewer than 250 detected genes, a library-complexity value of 0.80 or less, or a mitochondrial transcript fraction of 20% or greater were removed. Genes detected in fewer than 10 retained cells were subsequently excluded. Doublets were identified with DoubletFinder after SCTransform normalization, variable-feature selection, scaling, PCA, and UMAP analysis using principal components 1–10. The neighborhood parameter was evaluated by parameter sweep, and the recorded analysis used pN = 0.25 and pK = 0.09. The expected number of doublets was calculated using a 7.5% doublet rate and adjusted for the estimated homotypic doublet proportion. Only cells classified as singlets were retained.

Following quality control, cells were separated by sample and scored using S- and G2/M-phase marker sets. Each sample was normalized with SCTransform while regressing the mitochondrial transcript fraction. Datasets were integrated using the Seurat SCT workflow with 2,000 integration features, PrepSCTIntegration, FindIntegrationAnchors, and IntegrateData. PCA was performed on the integrated object, and UMAP and shared-nearest-neighbor graphs were generated using principal components 1–40. Graph-based clustering was assessed at resolutions 0.2, 0.4, 0.8, 1.2, 1.8, and 2.0; resolution 0.8 was used for downstream analysis. Cell identities were assigned according to normalized expression of canonical pancreatic cell-type markers and cross-checked by reference mapping to the Azimuth human pancreas reference and by anchor-based mapping to an internal pancreatic single-cell reference using FindTransferAnchors and MapQuery.

Single-cell gene-set enrichment analysis was performed using irGSEA on normalized RNA expression values. MSigDB human C2 gene sets were scored with AUCell, UCell, and singscore; C3 gene sets were scored with AUCell, UCell, singscore, and single-sample GSEA. Method-specific results were integrated by robust rank aggregation, and pathways were ordered by the robust-rank-aggregation P value. Differentially expressed genes were identified separately within each annotated cell type using Seurat FindMarkers on the RNA assay. Cytokine-treated cells were compared with control, exendin-4 plus cytokines, harmine plus cytokines, and harmine plus exendin-4 plus cytokines using a minimum detection fraction of 0.1 and an absolute log2 fold-change threshold of 0.25.

### Analysis of non-diabetic and T1D donor islet scRNA-seq data from the Human Pancreas Analysis Program (HPAP) database

To evaluate *SNHG6* and *TMA7* changes in β cells in T1D, we processed the publicly available scRNA-seq data from the Human Pancreas Analysis Program (HPAP) database (RRID:SCR_016202, https://hpap.pmacs.upenn.edu/) consortia under the Human Islet Research Network (HIRN) (RRID:SCR_014393)40 obtained from islets of 27 adult non-diabetic and 9 adult T1D cadaveric donors using CellRanger (V7.1.0) on the 10X cloud platform, referencing GRCh38-2020-A transcriptome.

### Human islet cell cultures and determination of β-cell death, inflammation marker gene expression, insulin secretion and HLA immunostaining

Human islet cells were cultured as previously reported (14) and incubated with proinflammatory cytokines, H, E or H+E for a period of 24 h and then fixed in 2% paraformaldehyde. In a different set of experiments, cells were treated for 24h with Thapsigargin (0.5µM), H, E and H+E. In an additional set of experiments, SNHG6 was knockdown with Adeno-U6-shRNA-SNHG6, or the expression enhanced with Adeno-CMV-SNHG6 (Vector Builder, Chicago, IL) and cells treated as above. β-Cell death was determined by the terminal deoxynucleotidyl transferase-mediated dUTP nick end-labeling (TUNEL) method (Promega, Madison, WI) and insulin (GN-ID4, Developmental Studies Hybridoma Bank, DSHB, Iowa City, IA) and DAPI co-immunolabeling. At least 2,000 β-cells per treatment were counted. Human islet cells were also immunolabeled for HLA-ABC BD (Pharmingen, Cat # 567855) or HLA-E (BD Pharmingen, Cat # 567855) together with insulin and DAPI co-immunostaining.

Analysis of *CASP3*, *iNOS*, *PSMB8, PSMB9, PSMB19, sXBP1, ATF3, TXNIP, ATF6, HLA-B, CXCL10, CHOP, SNHG6, IRF1* and *IRF2* mRNA expression in isolated islets or EndoC-βH1 cells was performed by real-time PCR using specific primers (**Supplementary Table 2**). Medium (100 μl) from islet and cell cultures was analyzed for nitric oxide (NO) by adding an equal volume of Greiss reagent (53). CXCL10 concentration in medium was determined using a specific ELISA (R&D Systems, Minneapolis, MN).

Analysis of insulin and proinsulin secretion analysis was performed in human islets following incubation in 2.8 or 16.8 mM glucose. Supernatants were collected, islets were washed with PBS and homogenized in acid/ethanol to measure total islet insulin content. Proinsulin and insulin levels were measured by ELISA (Mercodia, cat # 10-1118-01 and cat# 10-1113-01). Results were calculated as proinsulin or insulin secreted per insulin content and the ratio proinsulin/insulin was calculated.

### CyTOF Analysis

To precisely characterize cellular phenotypes at the single-cell level, we employed CyTOF (Cytometry by Time-of-Flight). Human peripheral blood mononuclear cells (PBMCs), provided by the Human Immune Monitoring Center at Icahn School of Medicine at Mount Sinai, from three different healthy subject donors were each divided into four aliquots (3x10^6^ cells/mL) and treated with vehicle (0.1% DMSO), H (10µM), E (10nM), or H+E in the presence of 1 µg/mL anti-CD3 (Biolegend, Cat#317347) plus 1 µg/mL anti-CD28 (Biolegend, cat#302943) plus IL-2 at 25 U/mL (Biolegend, cat#589106). Medium was changed every 3 days and after 12 days of treatment, cells were harvested and processed to perform CyTOF (Fluidigm Helios CyTOF 2 Mass Cytometer, Standard BioTools, Boston, MA) in the Human Immune Monitoring Center at Icahn School of Medicine using metal-conjugated antibodies to detect 16 immune cell type identification markers (*CD45, CD8, CD56, CD127, CD4, CD45RA, CD16, CD19, CD1c, CD66b, CD3, CD103, CD11b, CD11c, CD14,* and *CD123*) along with 22 functional, state markers and transcription factors [*Perforin, Granzyme B, Tbet, FoxP3, RORgT, EOMES, KLRG1, CD38, CD25, CD27, CD39, HLADR, TIM3, TOX, GATA3, PD-1, CD69, TIGIT, CCR7(CD197), TCF1, CCR6, 41BB(CD137) and CD57].* After we obtained sample data, panel data and flow cytometry standard files (fcs), we generated flowset data with flowcore package, then labeled the columns accordingly. We next structured the data into singleCellExperiment format, then embedded expression, log counts and metadata into the Seurat object for the convenience in the integration and visualization process. Subsequently, we applied centered log ratio (CLR) normalization and subset 300,000 cells to minimize the resource burden. Subsequently, the Seurat embedded data were integrated using reciprocal PCA (rPCA) reduction by assigning hyperparameter length of cell type identification markers. After running UMAP and neighboring algorithms, cell types were assigned using cell type markers in Louvain resolution of 0.4.

### NOD mice

Twelve-to-sixteen-week-old NOD/LtJ (NOD) female mice (The Jackson Laboratory) were housed in specific pathogen-free conditions. Non-fasting blood glucose was measured once a week by a portable glucometer (AlphaTRAK 2; Abbott Laboratories) until it reached ≥250 mg/dL when it was measured daily. Mice were considered diabetic when blood glucose was ≥250 mg/dL for three consecutive days and then iv treated once daily for 3 days with 5µg anti-CD3 antibody (non–Fc-binding monoclonal anti-CD3ε F(ab’)2 obtained from Bio X Cell, https://bxcell.com/product/m-CD3e-fab2-fragments/) or IgG (31). On the last day of iv treatment, an Alzet (Cupertino, CA) model 1004 mini-osmotic pump was subcutaneously implanted in each mouse to deliver at a continuous rate 3 mg/kg/day harmine, 0.1mg/kg/day exendin-4, vehicle (water) or the combination for 28 days. On day 28, pumps were replaced with new pumps containing fresh harmine and exendin-4 for additional 28 days of treatment. All procedures were performed with the approval of and in accordance with guidelines established by the Icahn School of Medicine at Mount Sinai and City of Hope Institutional Animal Care and Use Committees (IACUC #2015–0107 and IACUC #23030, respectively). Plasma insulin was determined by ELISA for mouse insulin (Mercodia). An intraperitoneal glucose tolerance test (IPGTT) was performed as described (14).

### Immunohistochemistry and Insulitis

At the end of the treatment, animals were sacrificed, pancreases harvested, fixed overnight at room temperature in 4% paraformaldehyde (PFA; Electron Microscopy Sciences, Hatfield, PA, USA), paraffin-embedded and analysis of β-cell mass, proliferation and death was performed in sections immunolabeled with anti-insulin (A5064, Agilent, Santa Clara, CA or 6N-ID4, Developmental Studies Hybridoma Bank, DSHB, Iowa City, IA) and anti-Ki67 (MA5-14520, Thermo Scientific, Waltham, MA) antibodies and DAPI, as previously described (14). Labeled cells were then visualized using fluorescent phase contrast microscopy (Zeiss Axiovert, White Plains, NY) and β-cell proliferation was quantified as Ki67+ and insulin+ and 1,000-2,000 β-cells blindly counted per sample. β-cell death was determined in sections stained for insulin using the TUNEL method (Promega, Madison, WI) (14) and at least 1,000 β-cells were blindly counted per sample. β-cell mass was measured in insulin-stained pancreas sections using ImageJ (National Institutes of Health, Bethesda, MD) (53). Sections were also stained with hematoxylin– eosin or insulin, DAPI and anti-CD45 (Novus Biologicals, cat# NB100-77417), anti-CD4 (SYSY, Cat# HS-360004), anti-CD8 (SYSY, Cat# HS-361003), anti-CD3 Abcam, (ab16669) or anti-B220 (Invitrogen, cat# 14045282) for pathologic evaluation of islet insulitis.

### Flow Cytometry Analysis

Spleen, pancreatic lymph nodes (PLNs) and pancreatic islets were harvested from NOD treated mice at the end of treatment and made into a cell suspension after lysis of the red blood cells and filtration. Cells (10^5^-10^6^ cells/mL) were treated for 16 h with 2 μg/mL soluble anti-CD3 and 2 μg/mL soluble anti-CD28 (BioLegend). Surface and intracellular staining of T cells for flow cytometry in the BD LSRFortessa™ Cell Analyzer (Waters, Milford MA) was achieved with FITC anti-mouse CD8b.2 Antibody (BioLegend, Cat# 140403); Brilliant Violet 421™ anti-mouse CD4 Antibody (BioLegend, Cat# 100437); PE anti-mouse IFN-γ Antibody (BioLegend, 505808); PE anti-mouse FOXP3 Recombinant Antibody (BioLegend, 118903). Live/dead cells were identified by the Zombie NIR Fixable Viability Kit (BioLegend).

### Statistical analysis

Data are presented as means ± SE in bar graphs, violin plots, scatterplots, and text. Statistical significance analysis was performed using Wilcoxon rank-sum test, two-tailed Student’s t-test or one-way ANOVA with Tukey’s multiple comparisons test for comparison among groups as indicated in the figure legends. P ≤ 0.05 was considered statistically significant. The simplified asterisk statistical significance annotation followed conventional criteria of 0.05, 0.01, 0.001, and 0.0001 for increment number of asterisks.

## Supplemental Figure Legends

**Supplemental Figure 1. Single cell RNA-seq analysis of human islets acutely treated with cytokines and harmine and exendin-4. A** Unsupervised clustering and cell type annotated UMAP with Louvain resolution 0.8 of non-diabetic human islets treated with vehicle, cytokines and harmine plus exendin-4. N=3 different human islet preparations. **B** Dot-plot visualization of top 4 canonical genes expressed in each cluster. **C** UMAP by individual treatment. **D** Cell type proportion in each annotated cluster. **E** Cell type proportion in each sample and treatment.

**Supplemental Figure 2. HLA-ABC and HLA-E immunolabeling of human islet cells treated with cytokines, harmine and exendin-4.** Representative images showing the immunolabeling for DAPI (blue), insulin (green), HLA-ABC (red) and HLA-E (magenta). Quantitation appears in Fig. 2, panels B and C.

**Supplemental Figure 3. Treatment of recent onset diabetic NOD mice with IgG and harmine and exendin-4. A** Random blood glucose levels in recent onset NOD mice treated iv for 3 days with IgG as control for anti-CD3 and then implanted with Alzet minipumps to deliver vehicle (H_2_O), harmine, exendin-4 or harmine+exendin-4 for 56 days. Dash line indicates 250mg/dL blood glucose. N=10 mice were used in each of the groups: IgG+H_2_O IgG+H, IgG+E, IgG+H+E. **B** Ratio of diabetic mice during the treatment period. **C** Non fasting plasma insulin levels at the end of treatment. One-way ANOVA with Tukey’s multiple comparisons test.

**Supplemental Figure 4. Presence of CD45+ cells in islets from NOD mice treated with anti-CD3+harmine+exendin-4. A** Schematic representation of the experiments. Created in BioRender. Garcia-Ocana, A. (2026). **B** Representative flow cytometry plots and quantification of CD45+ cells in total islet cells from islets isolated from αCD3- and H2O or H+E treated mice. N=3 mice per group. Blue=DAPI. Two-tailed Student’s t-test, *P<0.05.

**Supplemental Figure 5. CD45+ cells in spleen of NOD mice treated with anti-CD3+harmine+exendin-4. A** Representative flow cytometry plots depicting the immunophenotyping strategy. **B** Quantitation of CD45+ cells and the CD4/CD8 T-cell ratios in spleen of anti-CD3 (αCD3)-H_2_O- and αCD3-Harmine (H)+Exendin-4 (E)-treated mice at the end of treatment (day 56). N=10-12/group. Two-tailed Student’s t-test, *P<0.05.

**Supplemental Figure 6. CyTOF data analyzed using algorithm-based analysis. A** After embedding the CyTOF data into Seurat, we performed integration and visualization. UMAP shows PBMCs overlapping from the three different human PBMC donors. **B** UMAP shows PBMCs overlapping for the four different treatments which confirmed comparable distribution of cells across treatment groups. **C** Unsupervised clustering and cell type annotated UMAP with Louvain resolution 0.4 of PBMCs from non-diabetic subjects treated with vehicle (1% DMSO), harmine (H), exendin-4 (E) and H+E. N=3 different human PBMC preparations. **D** Dot-plot visualization of marker genes defining different immune populations and expressed in each cluster. **E** Dot-plot visualization of immune marker genes and annotated immune cell types. **F** UMAP shows major immune-cell populations assigned using established markers, including CD8+ T cells, CD4+ T cells, NK cells, B cells, and myeloid cells. Below separated by treatment.

**Supplemental Figure 7. SNHG6 Expression in EndoC-βH1 cells. A** Quantitation of *SNHG6* mRNA expression normalized by ACTIN expression assessed by qPCR in EndoC-βH1 cells following transduction with Adv-CMV or Adv-SNHG6-GPF. N=3, **P<0.01. **B** Quantitation of SNHG6 mRNA expression normalized by ACTIN expression assessed by qPCR in EndoC-βH1 cells following transduction with Adv-U6-sh-scrambled (SCR) or Adv-shSNHG6. N=3, Two-tailed Student’s t-test, **P<0.01.

## References

1. Atkinson MA, Eisenbarth GS, Michels AW. Type 1 diabetes. Lancet. 2014, 383(9911):69–82.

2. Quattrin T, Mastrandrea LD, Walker LSK. Type 1 diabetes. Lancet. 2023, 401(10394):2149–2162.

3. Herold KC, Delong T, Perdigoto AL, Biru N, Brusko TM, Walker LSK. The immunology of type 1 diabetes. Nat Rev Immunol 2024, 24:435–451.

4. Atkinson MA, Mirmira RG. The pathogenic “symphony” in type 1 diabetes: A disorder of the immune system, β cells, and exocrine pancreas. Cell Metab. 2023, 35(9):1500–1518.

5. Herold KC, Evans-Molina C. New and emerging therapies in type 1 diabetes mellitus. J Clin Invest. 2026, 136(10):e205520.

6. Mathieu C, Sims EK, Chatenoud L, James EA, Atkinson MA, Herold KC. Toward Disease-Modifying Therapies in Type 1 Diabetes: Focus on Teplizumab. Diabetes Care. 2026, 49(3):365–374.

7. Herold KC et al. An Anti-CD3 Antibody, Teplizumab, in Relatives at Risk for Type 1 Diabetes. N Engl J Med. 2019, 381:603–613.

8. Sims EK, Bundy BN, Stier K, Serti E, Lim N, Long SA, Geyer SM, Moran A, Greenbaum CJ, Evans-Molina C, Herold KC; Type 1 Diabetes TrialNet Study Group. Teplizumab improves and stabilizes beta cell function in antibody-positive high-risk individuals. Sci Transl Med. 2021, 13(583):eabc8980.

9. Ramos EL, Dayan CM, Chatenoud L, Sumnik Z, Simmons KM, Szypowska A, Gitelman SE, Knecht LA, Niemoeller E, Tian W, Herold KC; PROTECT Study Investigators. Teplizumab and β-Cell Function in Newly Diagnosed Type 1 Diabetes. N Engl J Med. 2023, 389(23):2151–2161.

10. Long SA, Thorpe J, Herold KC, Ehlers M, Sanda S, Lim N, Linsley PS, Nepom GT, Harris KM. Remodeling T cell compartments during anti-CD3 immunotherapy of type 1 diabetes. Cell Immunol. 2017, 319:3–9.

11. Penaranda C, Tang Q, Bluestone JA. Anti-CD3 therapy promotes tolerance by selectively depleting pathogenic cells while preserving regulatory T cells. J Immunol. 2011, 187(4):2015–22.

12. Wang P, Alvarez-Perez JC, Felsenfeld DP, Liu H, Sivendran S, Bender A, Kumar A, Sanchez R, Scott DK, Garcia-Ocaña A, Stewart AF. A high-throughput chemical screen reveals that harmine-mediated inhibition of DYRK1A increases human pancreatic beta cell replication. Nat Med. 2015 Apr;21(4):383–8.

13. Ackeifi C, Wang P, Karakose E, Manning Fox JE, González BJ, Liu H, Wilson J, Swartz E, Berrouet C, Li Y, Kumar K, MacDonald PE, Sanchez R, Thorens B, DeVita R, Homann D, Egli D, Scott DK, Garcia-Ocaña A, Stewart AF. GLP-1 receptor agonists synergize with DYRK1A inhibitors to potentiate functional human β cell regeneration. Sci Transl Med. 2020, 12(530):eaaw9996.

14. Rosselot C, Li Y, Wang P, Alvarsson A, Beliard K, Lu G, Kang R, Li R, Liu H, Gillespie V, Tzavaras N, Kumar K, DeVita RJ, Stewart AF, Stanley SA, Garcia-Ocaña A. Harmine and exendin-4 combination therapy safely expands human β cell mass in vivo in a mouse xenograft system. Sci Transl Med. 2024, 16(755):eadg3456.

15. Piganelli JD, Mamula MJ and James EA. The Role of β Cell Stress and Neo-Epitopes in the Immunopathology of Type 1 Diabetes. Front. Endocrinol. 2021, 11:624590.

16. Roep BO, Thomaidou S, van Tienhoven R, Zaldumbide A. Type 1 diabetes mellitus as a disease of the β-cell (do not blame the immune system?). Nat Rev Endocrinol. 2021, 17(3):150–161.

17. Eizirik, D.L., Miani, M. & Cardozo, A.K. Signalling danger: endoplasmic reticulum stress and the unfolded protein response in pancreatic islet inflammation. Diabetologia 2013, 56, 234–241.

18. Ramos-Rodríguez M, Raurell-Vila H, Colli ML, Alvelos MI, Subirana-Granés M, Juan-Mateu J, Norris R, Turatsinze JV, Nakayasu ES, Webb-Robertson BM, Inshaw JRJ, Marchetti P, Piemonti L, Esteller M, Todd JA, Metz TO, Eizirik DL, Pasquali L. The impact of proinflammatory cytokines on the β-cell regulatory landscape provides insights into the genetics of type 1 diabetes. Nat Genet. 2019 Nov;51(11):1588–1595.

19. Kulkarni A, Muralidharan C, May SC, Tersey SA, Mirmira RG. Inside the β Cell: Molecular Stress Response Pathways in Diabetes Pathogenesis. Endocrinology. 2022 Nov 14;164(1):bqac184.

20. Ponting, C.P.; Oliver, P.L.; Reik, W. Evolution and functions of long noncoding RNAs. Cell 2009, 136:629–641.

21. Mattick, J.S.; Amaral, P.P.; Carninci, P.; Carpenter, S.; Chang, H.Y.; Chen, L.L.; Chen, R.; Dean, C.; Dinger, M.E.; Fitzgerald, K.A.;, et al. Long non-coding RNAs: Definitions, functions, challenges and recommendations. Nat. Rev. Mol. Cell Biol. 2023, 24:430–447.

22. Long J, Badal SS, Ye Z, Wang Y, Ayanga BA, Galvan DL, et al. Long noncoding RNA Tug1 regulates mitochondrial bioenergetics in diabetic nephropathy. J Clin Invest (2016) 126(11):4205–18.

23. González-Moro I, Garcia-Etxebarria K, Mendoza LM, Fernández-Jiménez N, Mentxaka J, Olazagoitia-Garmendia A, et al. LncRNA ARGI Contributes to Virus-Induced Pancreatic β Cell Inflammation Through Transcriptional Activation of IFN-Stimulated Genes. Adv Sci (Weinh). 2023, 10(25):e2300063.

24. Gonzalez-Moro I, Olazagoitia-Garmendia A, Colli ML, Cobo-Vuilleumier N, Postler TS, Marselli L, et al. The T1D-associated lncRNA Lnc13 modulates human pancreatic β cell inflammation by allele-specific stabilization of STAT1 mRNA. Proc Natl Acad Sci U.S.A. 2020, 117(16):9022–31.

25. Wang Q, Wang Q, Meng X, Ji X, Wang T. LncRNA SNHG6 attenuates ferroptosis in high glucose-treated renal tubular epithelial cells by stabilizing YY1 to activate the PI3K/AKT/GSK-3β pathway. Arch Biochem Biophys. 2026, 776:110702.

26. Guo M, Dai Y, Jiang L, Gao J. Bioinformatics Analysis of the Mechanisms of Diabetic Nephropathy via Novel Biomarkers and Competing Endogenous RNA Network. Front Endocrinol (Lausanne). 2022, 13:934022.

27. Baldeón ME, Neece DJ, Nandi D, Monaco JJ, Gaskins HR. Interferon-gamma independently activates the MHC class I antigen processing pathway and diminishes glucose responsiveness in pancreatic beta-cell lines. Diabetes. 1997 46(5):770–8.

28. De George DJ, Ge T, Krishnamurthy B, Kay TWH, Thomas HE. Inflammation versus regulation: how interferon-gamma contributes to type 1 diabetes pathogenesis. Front Cell Dev Biol. 2023 11:1205590.

29. Khor B, Gagnon JD, Goel G, Roche MI, Conway KL, Tran K, Aldrich LN, Sundberg TB, Paterson AM, Mordecai S, Dombkowski D, Schirmer M, Tan PH, Bhan AK, Roychoudhuri R, Restifo NP, O’Shea JJ, Medoff BD, Shamji AF, Schreiber SL, Sharpe AH, Shaw SY, Xavier RJ. The kinase DYRK1A reciprocally regulates the differentiation of Th17 and regulatory T cells. Elife. 2015, 4:e05920.

30. Dandona P, Chaudhuri A, Ghanim H. Semaglutide in Early Type 1 Diabetes. N Engl J Med. 2023, 389(10):958–959.

31. Bresson D, Togher L, Rodrigo E, Chen Y, Bluestone JA, Herold KC, von Herrath M. Anti-CD3 and nasal proinsulin combination therapy enhances remission from recent-onset autoimmune diabetes by inducing Tregs. J Clin Invest. 2006, 116(5):1371–81.

32. Kaestner, K. H., Powers, A. C., Naji, A. & Atkinson, M. A. NIH initiative to improve understanding of the pancreas, islet, and autoimmunity in Type 1 diabetes: The Human Pancreas Analysis Program (HPAP). Diabetes 2019, 68, 1394–1402.

33. Xue S, Wasserfall CH, Parker M, Brusko TM, McGrail S, McGrail K, Moore M, Campbell-Thompson M, Schatz DA, Atkinson MA, Haller MJ. Exendin-4 therapy in NOD mice with new-onset diabetes increases regulatory T cell frequency. Ann N Y Acad Sci. 2008, 1150:152–6.

34. Sherry NA, Chen W, Kushner JA, Glandt M, Tang Q, Tsai S, Santamaria P, Bluestone JA, Brillantes AM, Herold KC. Exendin-4 improves reversal of diabetes in NOD mice treated with anti-CD3 monoclonal antibody by enhancing recovery of beta-cells. Endocrinology. 2007 Nov;148(11):5136–44.

35. Suarez-Pinzon WL, Power RF, Yan Y, Wasserfall C, Atkinson M, Rabinovitch A. Combination therapy with glucagon-like peptide-1 and gastrin restores normoglycemia in diabetic NOD mice. Diabetes. 2008, 57(12):3281–8.

36. Zumsteg U, Frigerio S, Holländer GA. Nitric oxide production and Fas surface expression mediate two independent pathways of cytokine-induced murine beta-cell damage. Diabetes. 2000, 49(1):39–47.

37. Arnush M, Heitmeier MR, Scarim AL, Marino MH, Manning PT, Corbett JA. IL-1 produced and released endogenously within human islets inhibits beta cell function. J Clin Invest. 1998, 102(3):516–26.

38. Eizirik DL, Pasquali L, Cnop M. Pancreatic β-cells in type 1 and type 2 diabetes mellitus: different pathways to failure. Nat Rev Endocrinol 2020; 16:349–362

39. Colli ML, Ramos-Rodríguez M, Nakayasu ES, Alvelos MI, Lopes M, Hill JLE, Turatsinze JV, Coomans de Brachène A, Russell MA, Raurell-Vila H, Castela A, Juan-Mateu J, Webb-Robertson BM, Krogvold L, Dahl-Jorgensen K, Marselli L, Marchetti P, Richardson SJ, Morgan NG, Metz TO, Pasquali L, Eizirik DL. An integrated multi-omics approach identifies the landscape of interferon-α-mediated responses of human pancreatic beta cells. Nat Commun. 2020, 11(1):2584.

40. Frigerio S, Junt T, Lu B, Gerard C, Zumsteg U, Holländer GA, Piali L. Beta cells are responsible for CXCR3-mediated T-cell infiltration in insulitis. Nat Med. 2002, 8(12):1414–20.

41. Sandor AM, Jacobelli J, Friedman RS. Immune cell trafficking to the islets during type 1 diabetes. Clin Exp Immunol. 2019, 198(3):314–325.

42. Engin F, Yermalovich A, Nguyen T, Hummasti S, Fu W, Eizirik DL, Mathis D, Hotamisligil GS. Restoration of the unfolded protein response in pancreatic β cells protects mice against type 1 diabetes. Sci Transl Med. 2013, 5(211):211ra156.

43. Tersey SA, Nishiki Y, Templin AT, Cabrera SM, Stull ND, Colvin SC, Evans-Molina C, Rickus JL, Maier B, Mirmira RG. Islet β-cell endoplasmic reticulum stress precedes the onset of type 1 diabetes in the nonobese diabetic mouse model. Diabetes. 2012, 61(4):818–27.

44. Webster KL, Mirmira RG. Beta cell dedifferentiation in type 1 diabetes: sacrificing function for survival? Front Endocrinol (Lausanne). 2024, 15:1427723.

45. Emily K. Sims, Farooq Syed, Julius Nyalwidhe, Henry T. Bahnson, Leena Haataja, Cate Speake, Margaret A. Morris, Appakalai N. Balamurugan, Raghavendra G. Mirmira, Jerry Nadler, Teresa L. Mastracci, Peter Arvan, Carla J. Greenbaum, Carmella Evans-Molina. Abnormalities in proinsulin processing in islets from individuals with longstanding T1D. Translational Research, 2019, 213: 90–99.

46. Sims EK, Bahnson HT, Nyalwidhe J, Haataja L, Davis AK, Speake C, DiMeglio LA, Blum J, Morris MA, Mirmira RG, Nadler J, Mastracci TL, Marcovina S, Qian WJ, Yi L, Swensen AC, Yip-Schneider M, Schmidt CM, Considine RV, Arvan P, Greenbaum CJ, Evans-Molina C; T1D Exchange Residual C-peptide Study Group. Proinsulin Secretion Is a Persistent Feature of Type 1 Diabetes. Diabetes Care. 2019, 42(2):258–264.

47. Zheng X, Lu J, Qiu H, Jin Q, Cao L, Liu J, Song P, Yao S. SNHG6 Promotes Lipid Metabolic Reprogramming in Hepatocellular Carcinoma via Upregulation of SCD. J Hepatocell Carcinoma. 2026, 13:610313.

48. Zhang W, Zhou Y, Ye L, Huang C, Wang Y. Long non-coding RNA SNHG6 promotes odontoblastic differentiation of human dental pulp stem cells via the PI3K/Akt/mTOR pathway. Differentiation. 2026, 147:100927.

49. Liu F, Tian T, Zhang Z, Xie S, Yang J, Zhu L, Wang W, Shi C, Sang L, Guo K, Yang Z, Qu L, Liu X, Liu J, Yan Q, Ju HQ, Wang W, Piao HL, Shao J, Zhou T, Lin A. Long non-coding RNA SNHG6 couples cholesterol sensing with mTORC1 activation in hepatocellular carcinoma. Nat Metab. 2022, 4(8):1022–1040.

50. Chen D, Thayer TC, Wen L, Wong FS. Mouse Models of Autoimmune Diabetes: The Nonobese Diabetic (NOD) Mouse. Methods Mol Biol. 2020, 2128:87–92.

51. Thayer TC, Wilson SB, Mathews CE. Use of nonobese diabetic mice to understand human type 1 diabetes. Endocrinol Metab Clin North Am. 2010, 39(3):541–61.

52. Kumar K, Wang P, Wilson J, Zlatanic V, Berrouet C, Khamrui S, Secor C, Swartz EA, Lazarus M, Sanchez R, Stewart AF, Garcia-Ocana A, DeVita RJ. Synthesis and Biological Validation of a Harmine-Based, Central Nervous System (CNS)-Avoidant, Selective, Human β-Cell Regenerative Dual-Specificity Tyrosine Phosphorylation-Regulated Kinase A (DYRK1A) Inhibitor. J Med Chem. 2020, 63(6):2986–3003.

53. Lu G, Rausell-Palamos F, Zhang J, Zheng Z, Zhang T, Valle S, Rosselot C, Berrouet C, Conde P, Spindler MP, Graham JG, Homann D, Garcia-Ocaña A. Dextran Sulfate Protects Pancreatic β-Cells, Reduces Autoimmunity, and Ameliorates Type 1 Diabetes. Diabetes. 2020, 69(8):1692–1707.

