## Supplementary figures and images for "Harmine Plus Exendin-4 Enhances Remission of Recent-Onset Type 1 Diabetes Following Anti-CD3 Therapy"

### Supplemental Tables

**Supplemental Table 1.**

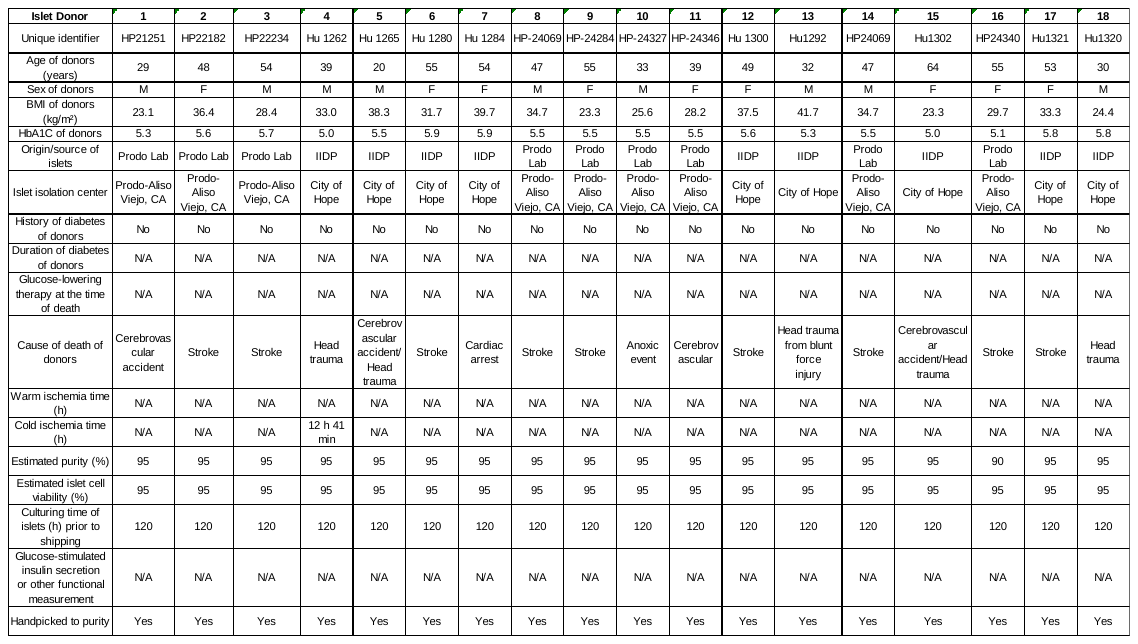

**Supplemental Table 2.**

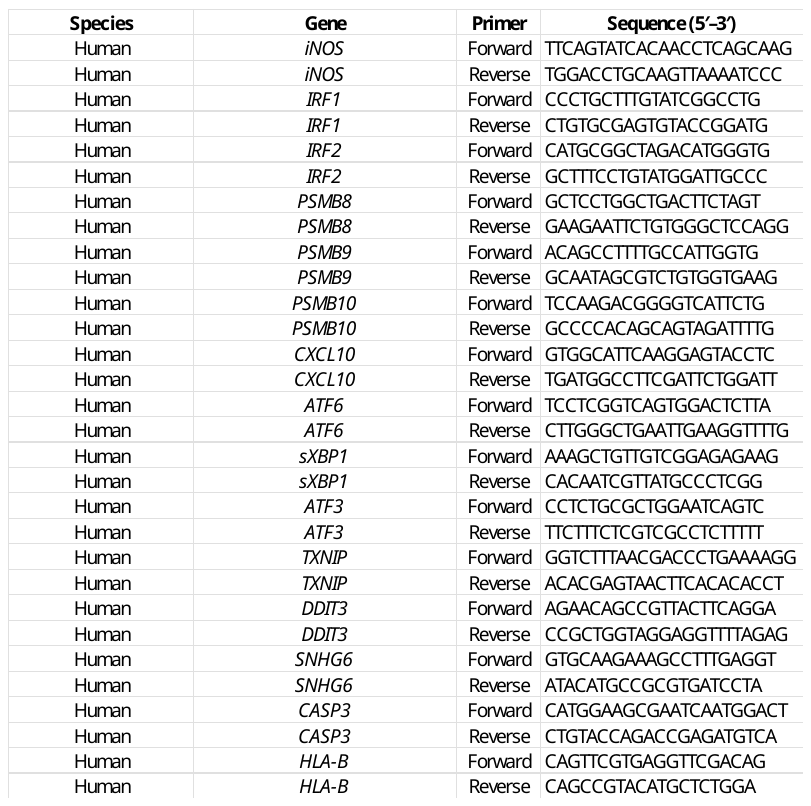
