## Supplemental Figures for "Harmine Plus Exendin-4 Enhances Remission of Recent-Onset Type 1 Diabetes Following Anti-CD3 Therapy"

### Supplemental Figure 1

A

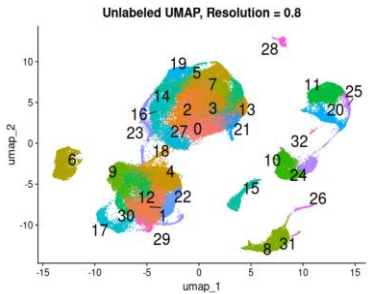

B

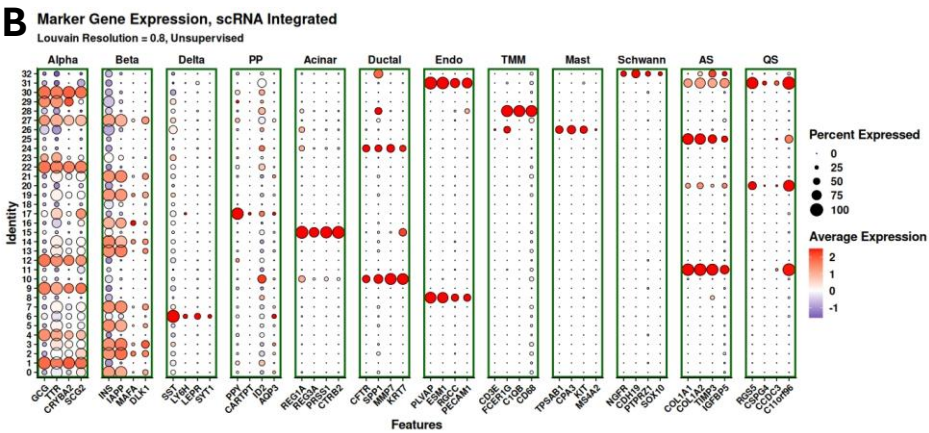

C

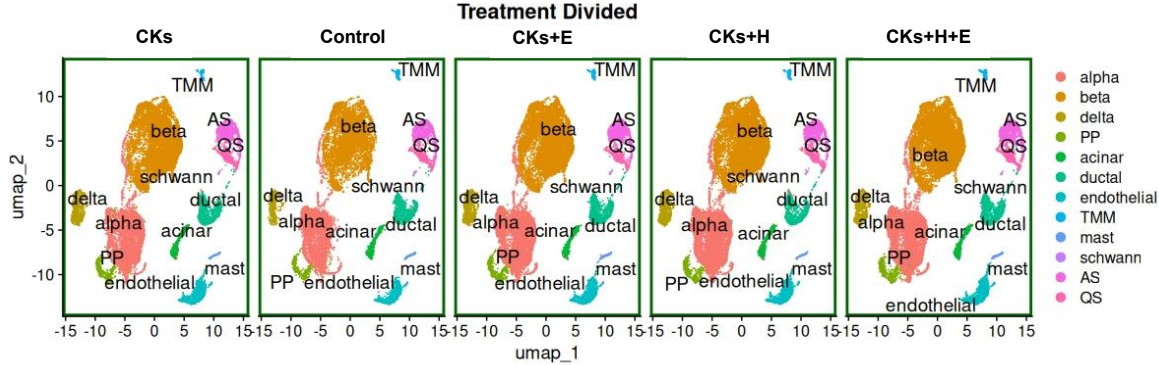

D

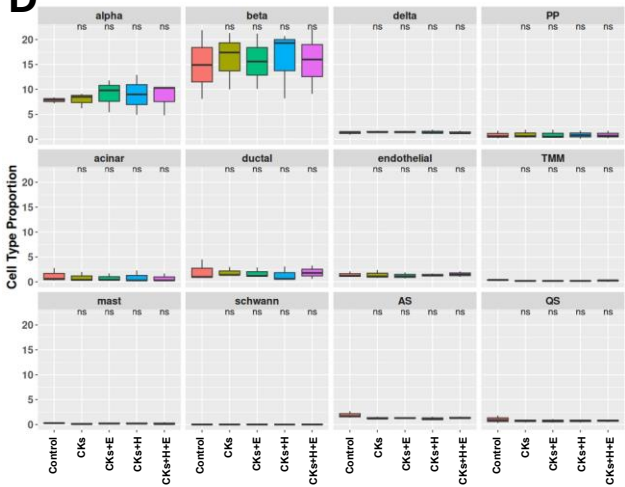

E

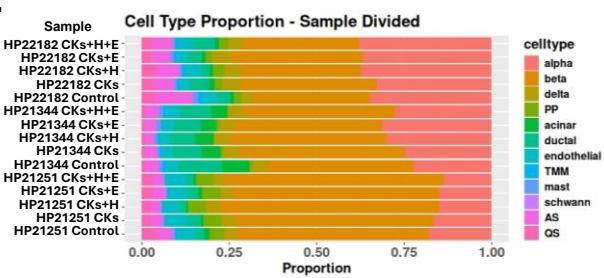

Supplemental Figure 2

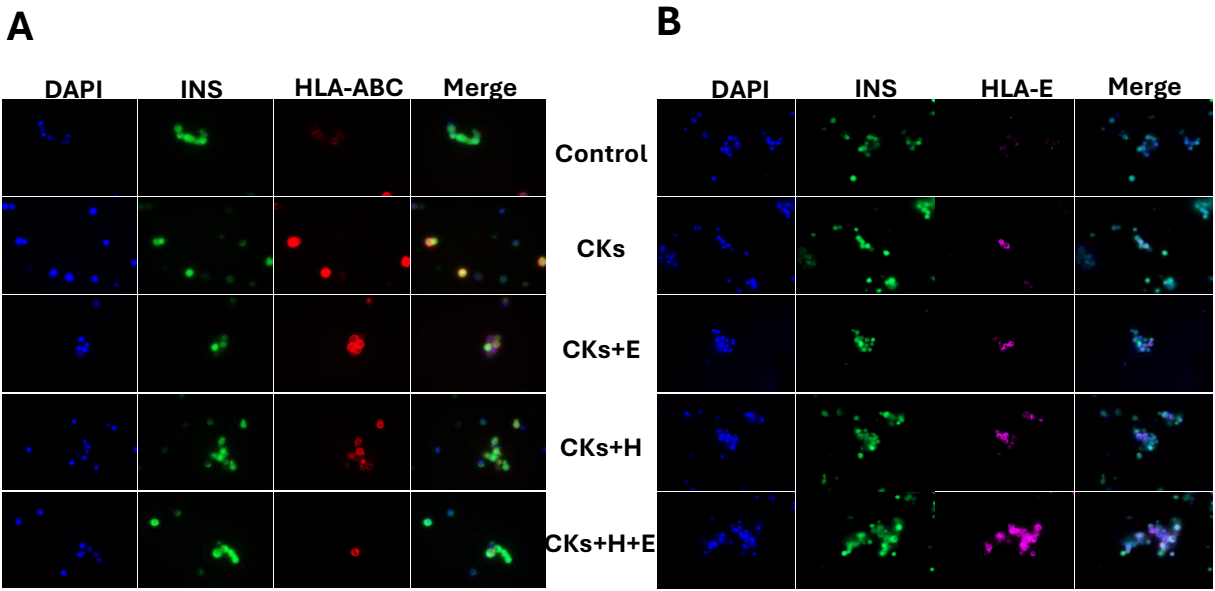

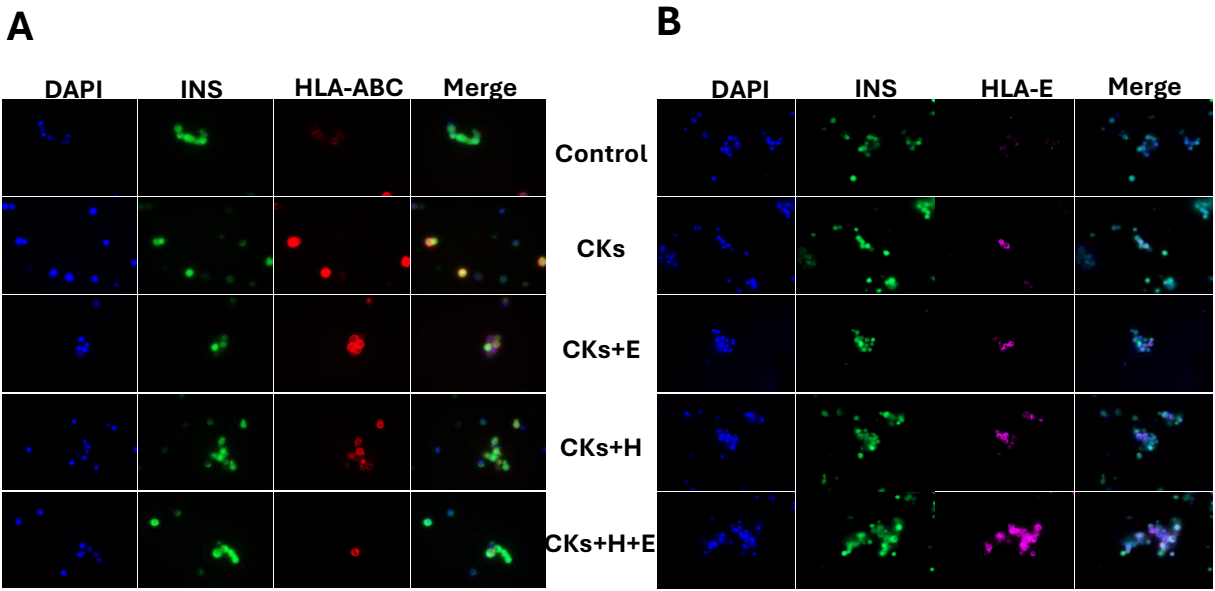

### Supplemental Figure 3

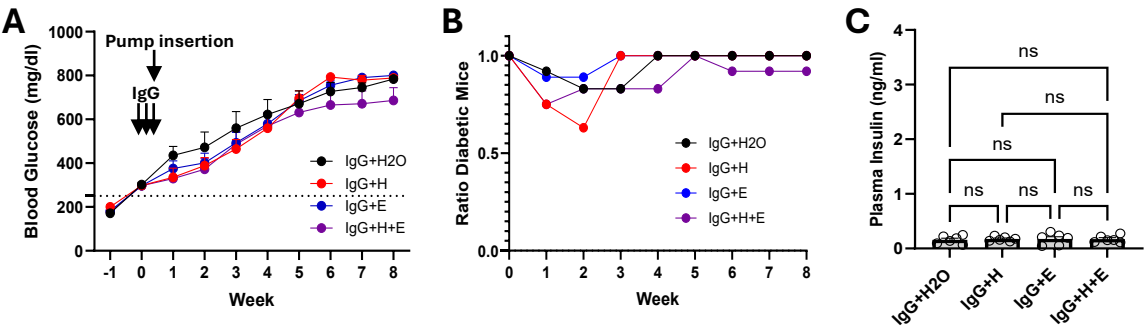

### Supplemental Figure 4

**A**

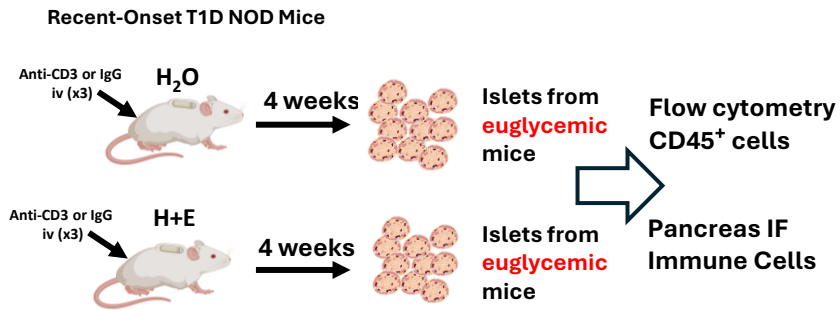

**B**

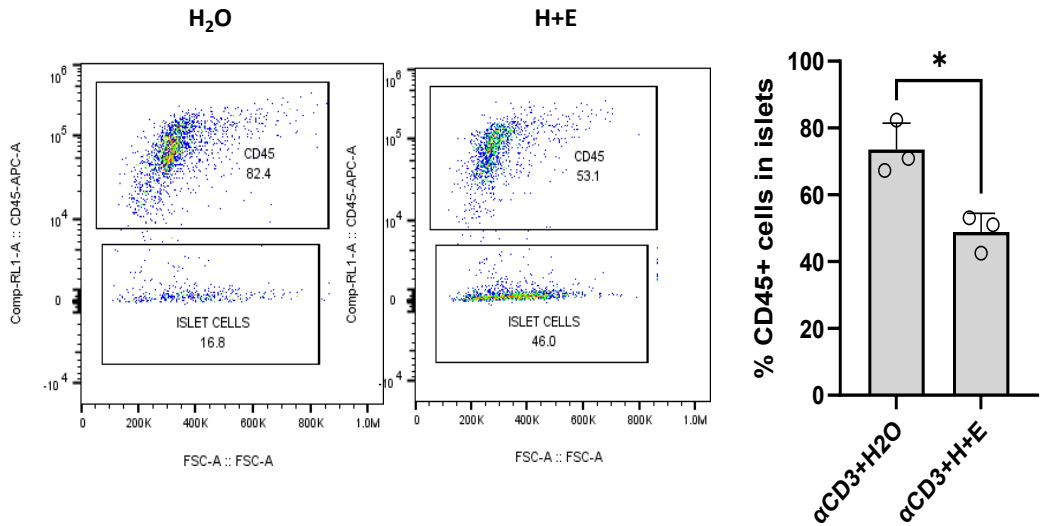

### Supplemental Figure 5

A

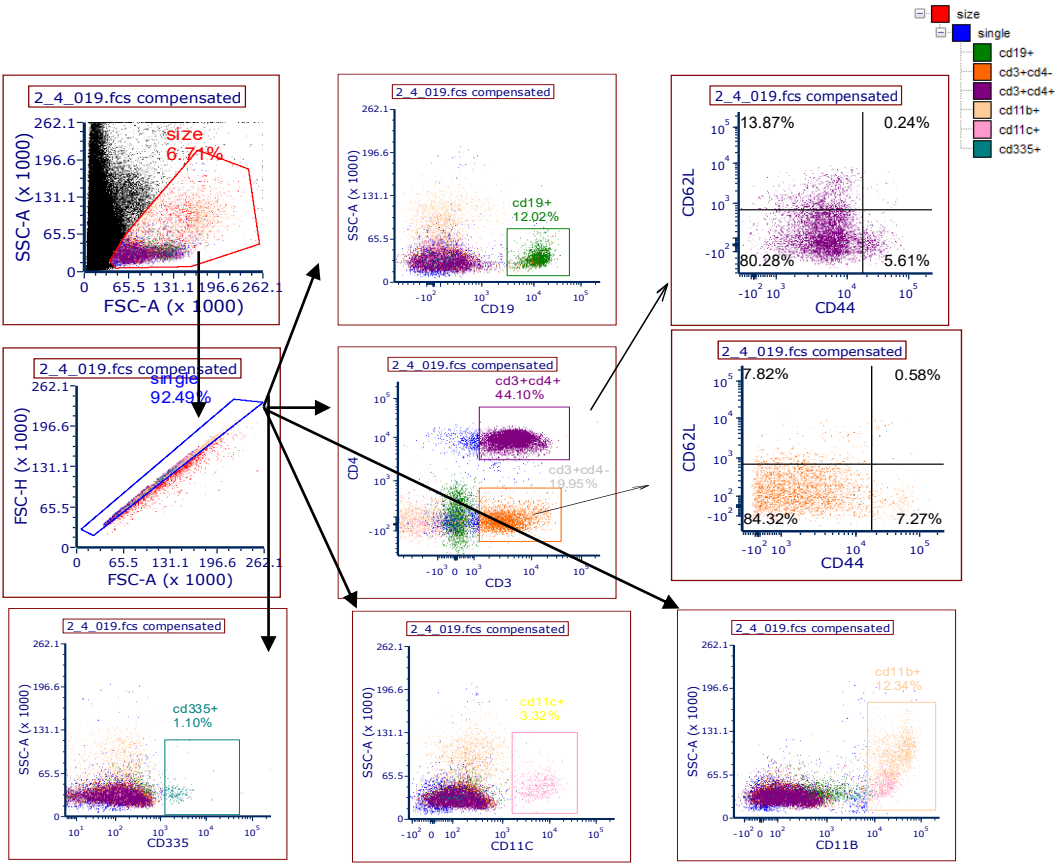

B

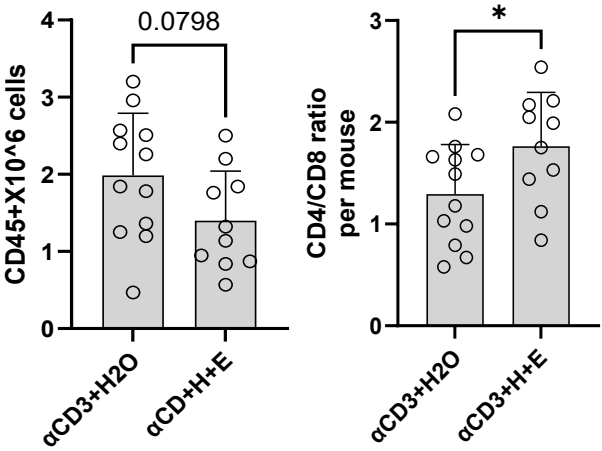

Supplemental Figure 6

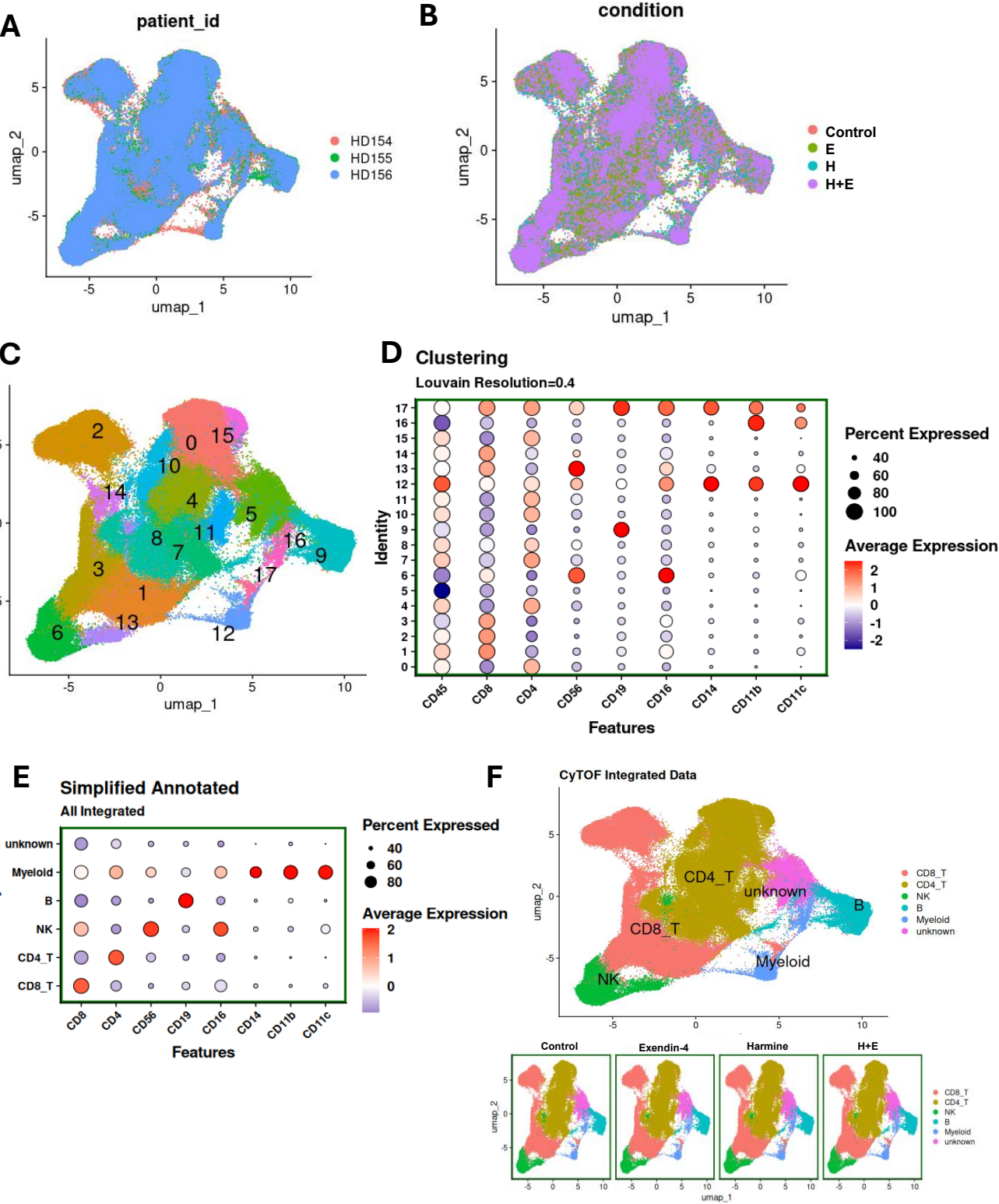

Supplemental Figure 7

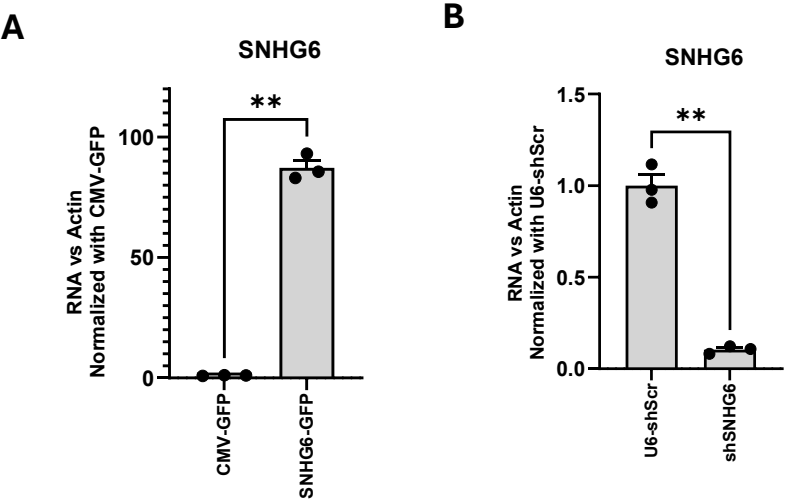
